# Lysosome repair by ER-mediated cholesterol transfer

**DOI:** 10.1101/2022.09.26.509457

**Authors:** Maja Radulovic, Eva Maria Wenzel, Sania Gilani, Lya K.K. Holland, Alf Håkon Lystad, Santosh Phuyal, Vesa M. Olkkonen, Andreas Brech, Marja Jäättelä, Kenji Maeda, Camilla Raiborg, Harald Stenmark

**Affiliations:** Centre for Cancer Cell Reprogramming, Faculty of Medicine, University of Oslo, Montebello, N-0379 Oslo, Norway; Department of Molecular Cell Biology, Institute for Cancer Research, Oslo University Hospital, Montebello, N-0379 Oslo, Norway; Cell Death and Metabolism, Center for Autophagy, Recycling and Disease, Danish Cancer Society Research Center, 2100 Copenhagen, Denmark; Department of Molecular Medicine, Institute of Basic Medical Sciences, University of Oslo, 0317 Oslo, Norway; Minerva Foundation Institute for Medical Research, Biomedicum 2U, Tukholmankatu 8, 00290, Helsinki, Finland; Department of Anatomy, Faculty of Medicine, 00014 University of Helsinki, Helsinki, Finland; Department of Cellular and Molecular Medicine, Faculty of Health Sciences, University of Copenhagen, DK-2200 Copenhagen, Denmark

## Abstract

Lysosome integrity is essential for cell viability, and lesions in lysosome membranes are repaired by the ESCRT machinery. Here we describe an additional mechanism for lysosome repair that is activated independently of ESCRT recruitment. Lipidomic analyses showed increases in lysosomal phosphatidylserine and cholesterol after damage. Electron microscopy demonstrated that lysosomal membrane damage is rapidly followed by formation of contacts with the endoplasmic reticulum (ER), which depend on the ER proteins VAPA/B. The cholesterol-binding protein ORP1L was recruited to damaged lysosomes, accompanied by cholesterol accumulation by a mechanism that required VAP-ORP1L interactions. The PtdIns 4-kinase PI4K2A rapidly produced PtdIns4P on lysosomes upon damage, and knockout of PI4K2A inhibited damage-induced accumulation of ORP1L and cholesterol and led to failure of lysosomal membrane repair. The cholesterol-PtdIns4P transporter OSBP was also recruited upon damage, and its depletion caused lysosomal accumulation of PtdIns4P and resulted in cell death. We conclude that ER contacts are activated on damaged lysosomes in parallel to ESCRTs to provide lipids for membrane repair, and that PtdIns4P generation and removal are central in this response.

## INTRODUCTION

Lysosomes are the main degradative organelles of the cell and play crucial roles in metabolism, signalling, homeostasis, and immunity (Ballabio & Bonifacino, 2020; Holland *et al*, 2020). Because of their high luminal concentrations of protons, Ca^2+^ and hydrolases, lysosome rupture can cause cell lethality through several pathways (Papadopoulos & Meyer, 2017). To prevent untimely death, cells are therefore equipped with mechanisms that repair damaged lysosomes, or degrade them if damage is beyond repair. The endosomal sorting complex required for transport (ESCRT) machinery is rapidly recruited to damaged lysosome membranes to mediate repair (Lopez-Jimenez *et al*, 2018; Radulovic *et al*, 2018; Skowyra *et al*, 2018), whereas the autophagy machinery is recruited with slower kinetics to sequester and degrade severely damage lysosomes (Hung *et al*, 2013; Maejima *et al*, 2013).

The observation that ESCRT depletion does not fully prevent lysosome repair has suggested that additional repair mechanisms may exist (Radulovic *et al*., 2018; Skowyra *et al*., 2018). Indeed, an ESCRT-independent mechanism that involves lysosomal sphingomyelin scrambling and consequent sphingomyelinase-mediated generation of ceramide has recently been described (Niekamp *et al*, 2022).

Here we have performed lipidomic analyses of isolated lysosomes at different time points after membrane damage to obtain a detailed view of lysosomal lipid composition upon damage. Motivated by our discoveries that lysosome damage causes increases in lysosomal phosphatidylserine (PS), cholesterol and a phosphorylated PtdIns species, accompanied by rapid mobilization of endoplasmic reticulum (ER) around damaged lysosomes, we have here uncovered a third mechanism for lysosome repair, executed via ER-lysosome contacts.

## RESULTS

### Changes in lysosome lipid composition upon membrane damage

To elucidate the effects of membrane damage on the lipid compositions of lysosomes, we quantitatively profiled lipids in HeLa cells treated with a lysosomotropic compound, L-leucyl-L-leucine methyl ester (LLOMe) (Thiele & Lipsky, 1990) for 0, 10, or 45 min. We immuno-affinity purified lysosomes from the prepared post-nuclear fractions using a primary antibody against LAMP1 and a secondary antibody conjugated to magnetic microbeads (Bilgin *et al*, 2017; Stahl-Meyer *et al*, 2022) (Fig EV1A).

Quantitative mass spectrometry-based shotgun lipidomics analysis of the purified lysosomes (LAMP1+ fraction) identified more than 200 species in 25 lipid classes belonging to the categories of glycerophospholipids (GPLs), sphingolipids (SLs), and sterol lipids (STs), and roughly the same numbers in the corresponding whole cell lysates (WCLs) (Figs 1B and EV1B). The purification yielded in total ∼1,500 picomoles (pmol) of these lipids from untreated (0 min) cells, in comparison to only ∼150 pmol in the negative control, in which the anti-LAMP1 antibody was replaced with a control IgG antibody (Figs 1A and EV1C). Further supporting its purity, the LAMP1+ fraction had in average ∼30 fold higher molar percentage (mol%, molar quantity relative to the total pmol of all lipids) values of bismonophosphatidic acid (BMP) species, the primarily polyunsaturated GPLs residing in the intraluminal vesicles of lysosomes (Nielsen *et al*, 2020; Stahl-Meyer *et al*., 2022; van Meer *et al*, 2008), than the WCL (Bilgin *et al*., 2017; Nielsen *et al*., 2020) (Fig EV1D,E). The LAMP1+ fraction was abundant in STs and SLs (Fig 1B) but depleted of most other diacyl-GPL classes than BMP, including the major lipid class of HeLa ER membrane phosphatidylcholine (PC) (Scrima *et al*, 2022) and the fully depleted mitochondrial cardiolipin (CL) (Figs EV1D and EV2).

**Figure 1.**
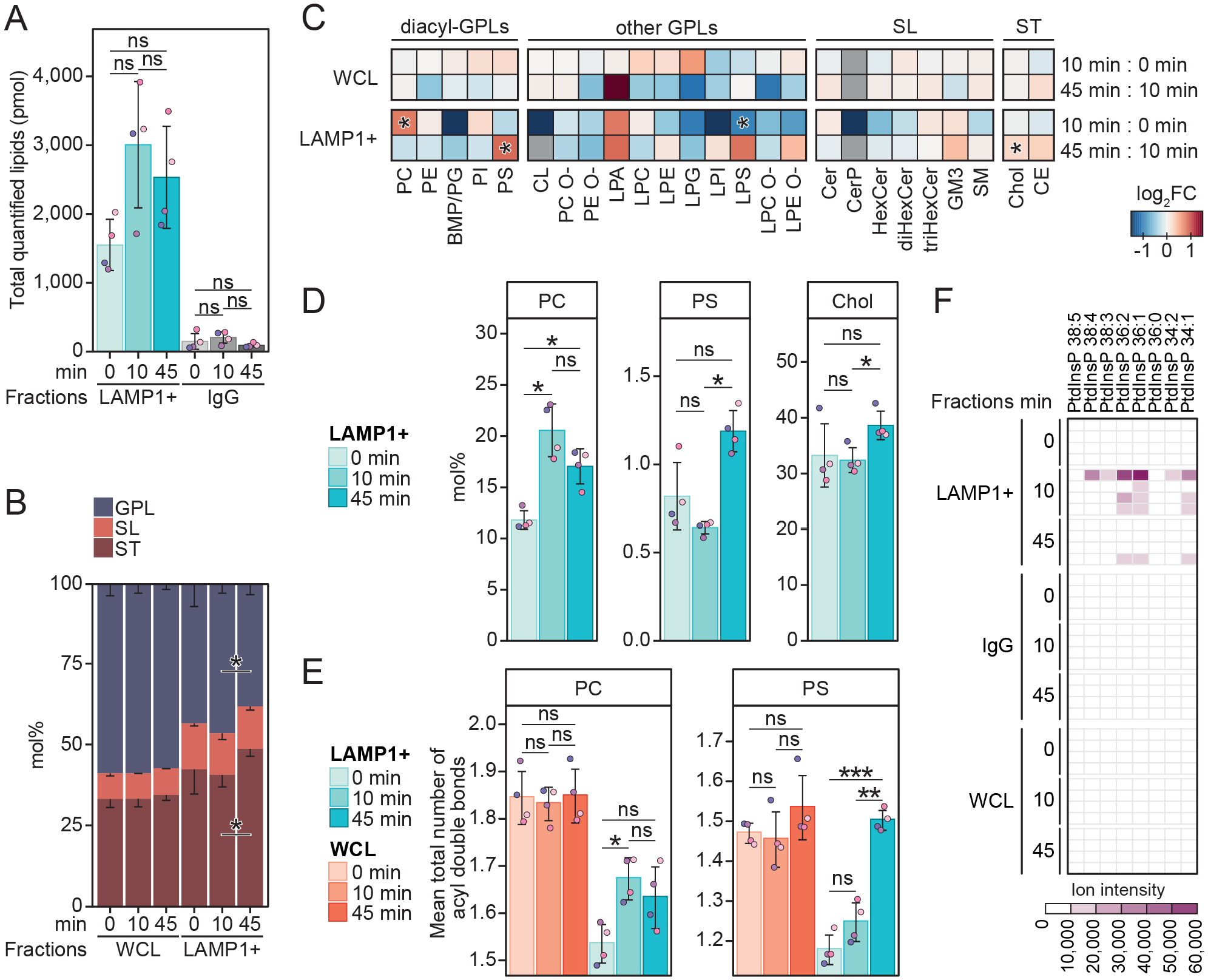
Lysosome lipid composition is altered upon membrane damage. **A** Total molar quantity of lipids identified in the produced LAMP1-positive (LAMP1+) and the corresponding IgG negative control fractions, produced from HeLa cells treated with LLOMe for 0, 10, or 45 min. **B** Molar percentages (mol%) values of lipid categories glycerophospholipids (GPL), sphingolipids (SL), and sterol lipids (ST) in the LAMP1+ fractions and the corresponding whole cell lysate (WCL). **C** LLOMe-induced fold change (log_2_-transformed) in the mol% values of lipid classes in the WCLs and LAMP1+ fractions, in the first 10 min (mol% at 10 min: mol% at 0 min) and the following 45 min (mol% at 45 min: mol% at 10 min) of treatment. **D** Mol% values of diacyl-GPL lipid classes phosphatidylcholine (PC) and phosphatidylserine (PS), and cholesterol (Chol) in the LAMP1+ fractions. **E** Mean total numbers of acyl double bonds in the PC and PS classes in the WCLs and LAMP1+ fractions. **F** Ion intensities of dehydrated phosphorylinositol headgroup recorded in MS2 after fragmentation of [M-2H]^2-^ ions of phosphatidylinositol phosphate (PtdInsP) species detected in the negative ion mode MS1. Intensities measured in the four replicates are displayed in individual rows. N=3-4. Multiple t-tests were performed to assess changes between means of pmol (A), lipid quantities in mol% (B, C, D), or the mean total numbers of double bonds in diacyl-GPL classes (E). Resultant *p*-values were corrected for multiple testing using Benjamini-Hochberg correction and differences were considered statistically significantly changed with adjusted *p*-values (q-values) of 0.05 or less. * q < 0.05; ** q < 0.01; *** q < 0.001;**** q < 0.0001. Heatmaps are for simplicity only fitted with a single asterisk regardless of the magnitude of the *p*-value. *Additional abbreviations*: BMP, bismonophosphatidic acid; CE, cholesteryl ester; Cer, ceramide; CerP, ceramide-1-phosphate; CL, cardiolipin; diHexCer, dihexosylceramide; GM3, ganglioside GM3; HexCer, hexosylceramide; LPA, lyso-phosphatidic acid; LPC, lysoPC; LPC O-, alkyl-LPC; LPE, lysoPE; LPE O-, alkyl-LPE; LPG, lysoPG; LPI, lysoPI; LPS, lysoPS; PC O-, acyl-alkyl-PC; PE, phosphatidylethanolamine, PE O-, acyl-alkyl-PE; PG, phosphatidylglycerol; PI, phosphatidylinositol; SM, sphingomyelin; triHexCer, trihexosylceramide.

LLOMe treatment for 10 min did not significantly alter the mol% values of the 25 monitored lipid classes in the WCL (Figs 1C and EV2). In contrast, this treatment already increased the total lipid amount yielded into the LAMP1+ fraction (to ∼ 3,000 pmol, not statistically significant) (Fig 1A). Moreover, it increased the PC class from 11.8 mol% to 20.5 mol% (Fig 1C, D) and rendered it less saturated (Figs 1E and EV3) to better resemble the PC class in the ER (Scrima *et al*., 2022; van Meer *et al*., 2008). LLOMe at 10 min did not significantly affect the mol% of any other lipid classes except the minor lysophosphatidylserine (LPS) (Fig 1C). Extending LLOMe treatment to 45 min did not enhance these trends observed after 10 min (Fig 1A,C), but instead increased the mol% values of cholesterol and the PS class (Fig 1C,D), of which the latter similarly to the PC class increased the mean total number of acyl double bonds (Figs 1E and EV3).

Cholesterol and the PS class are two specific ligands of OSBP-related proteins (ORPs) exporting these lipids out of the ER to late secretory compartments (Maeda *et al*, 2013; Moser von Filseck *et al*, 2015). The observation that membrane damage increased the lysosomal contents of these two lipids (Fig 1C,D) encouraged us to re-examine the shotgun lipidomics data for phosphatidylinositol 4-phoshate (PtdIns4P), yet another ligand of ORPs (Moser von Filseck *et al*., 2015). Even though we were unable to quantify them due to the lack of a PtdIns4P standard in our lipidomics pipeline, we detected precursor ions and associated fragment ions matching multiple PtdIns4P (or other PtdInsPs) species (Figs 1F and EV1G). These PtdInsP species were nearly exclusively detected in the LAMP1+ fraction after LLOMe treatment for 10 min, and not in any WCLs. Overall, the lipidomics data revealed that membrane damage rapidly increases PC and PtdInsP associated with lysosomes, followed by later elevations in cholesterol and the PS class.

### Lysosome damage induces VAP-dependent ER-lysosome contacts

The increase in lipid yield observed in the LAMP1+ fraction after LLOMe treatment suggests that lipid transfer is involved in lysosome repair. Because ER membranes are involved in lipid transfer (Ikonen & Zhou, 2021; Lagace & Ridgway, 2013), we investigated whether lysosome damage increases the extent of ER-lysosome contacts. Indeed, incubation of HeLa cells with LLOMe for only 5 minutes led to a strong increase in contact sites between ER and lysosomes, which was not further increased by more prolonged incubation (Fig 2A,B). Many of the known contact sites between ER and other organelles contain ER proteins of the VAP family, and we therefore monitored LLOMe-induced contacts in HeLa cells with double knockout of VAP-A and VAP-B (Dong *et al*, 2016). Interestingly, these cells showed no increase in ER-lysosome contacts upon LLOMe treatment (Fig EV4), indicating that the damage-induced ER-lysosome contacts are VAP dependent.

**Figure 2.**
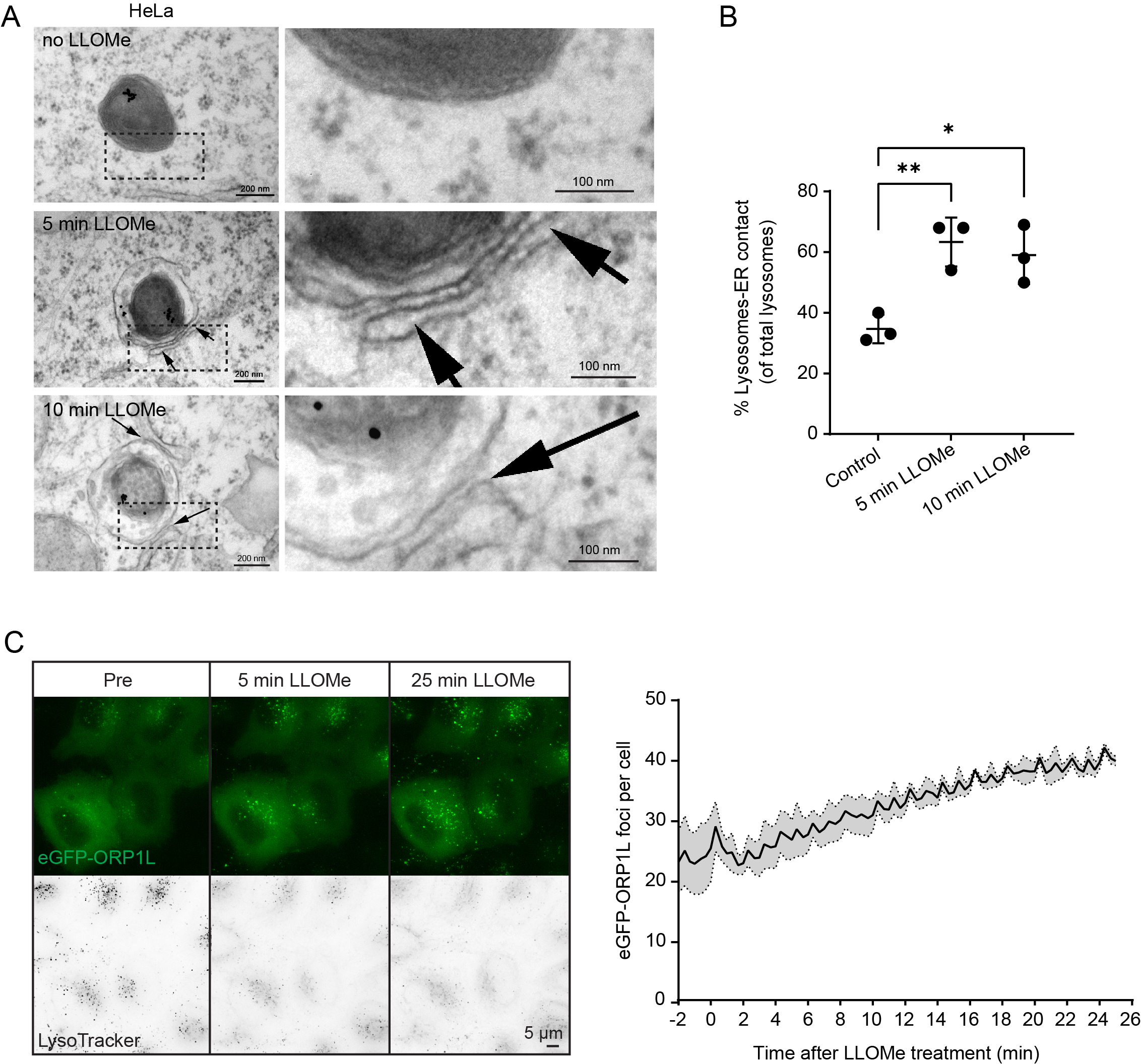
Lysosome damage induces VAP-dependent ER-Lysosome contacts. **A** Representative electron micrographs of HeLa cells treated or not with 250 µM LLOMe for 5 min or 10 min before chemical fixation. Cells were incubated for 4 h following washing steps and overnight incubation with 15 nm gold nanoparticles conjugated to bovine serum albumin to visualize lysosomes (black spots inside lysosomes are internalized gold nanoparticles) before treatment with LLOMe and processing samples for electron microscopy. Arrows indicate areas with increased lysosomes-endoplasmic reticulum contacts. **B** Quantification of electron micrographs showing an increase in membrane contact sites between ER and Lysosomes following 250 µM LLOMe treatment for 5 or 10 minutes. Error bars denote +/-SD from n=3 independent experiments, >48 lysosomes were quantified per experiment for each condition. Statistical significance was determined using one-way ANOVA with Dunnett’s multiple comparisons test, **** p<0.0001 **C** Representative movie stills of a live-cell imaging experiment illustrating the recruitment of transiently expressed eGFP-ORP1L to lysosomes following 250 µM LLOMe treatment. Cells were pre-treated for 30 min with 75 nM Lysotracker Deep Red to monitor lysosome damage (judged by the decreased number of Lysotracker spots after LLOMe treatment). The graph represents quantification of eGFP-ORP1L foci per cell after LLOMe treatment. Error bars denote +/-SEM from n=3 independent live-cell imaging experiments, >25 cells were analyzed per experiment.

### ORP1L is recruited to lysosomes upon damage

ORP family proteins constitute a group of proteins that form VAP-dependent contact sites between ER and other organelles (Olkkonen & Ikonen, 2022). Whereas several ORP proteins are involved in PS transport, others mediate cholesterol transport. Since the cholesterol transport protein ORP1L has been implicated in contacts between the ER and late endosomes/lysosomes (Rocha *et al*, 2009), we investigated whether this protein is recruited to lysosomes upon damage. Interestingly, live microscopy of HeLa cells expressing eGFP-tagged ORP1L showed that this protein is recruited to lysosomes already at 5 minutes after LLOMe addition, and recruitment was further increased up to 30 minutes of LLOMe (Fig 2C, Movie EV1). This suggests that ORP1L-containing ER-lysosome contacts are formed upon lysosome damage.

### ORP1L is recruited by PI4K2A-dependent accumulation of PtdIns4P on damaged lysosomes

The damage-induced recruitment of ORP1L to lysosomes raised the question of how such recruitment is triggered. Since ORP1L contains a phosphoinositide-binding PH domain (Boutry & Kim, 2021; Dong *et al*, 2019), and since the lipidomics data indicated rapid increase in a PtdInsP species after LLOMe, we examined cells expressing fluorescently tagged probes for various phosphoinositides and monitored their possible recruitment to lysosomes upon LLOMe treatment. Strikingly, PtdIns4P, visualized with a tandem PtdIns4P-binding domain from the bacterial protein SidM (Hammond *et al*, 2014), was rapidly recruited to lysosomes after LLOMe treatment (Fig 3A, Movie EV2), whereas probes for PtdIns3P, PtdIns(3,5)P_2_ and PtdIns(4,5)P_2_ showed no increase (Fig EV5). This indicates that PtdIns4P is acutely produced on lysosome membranes upon their damage.

**Figure 3.**
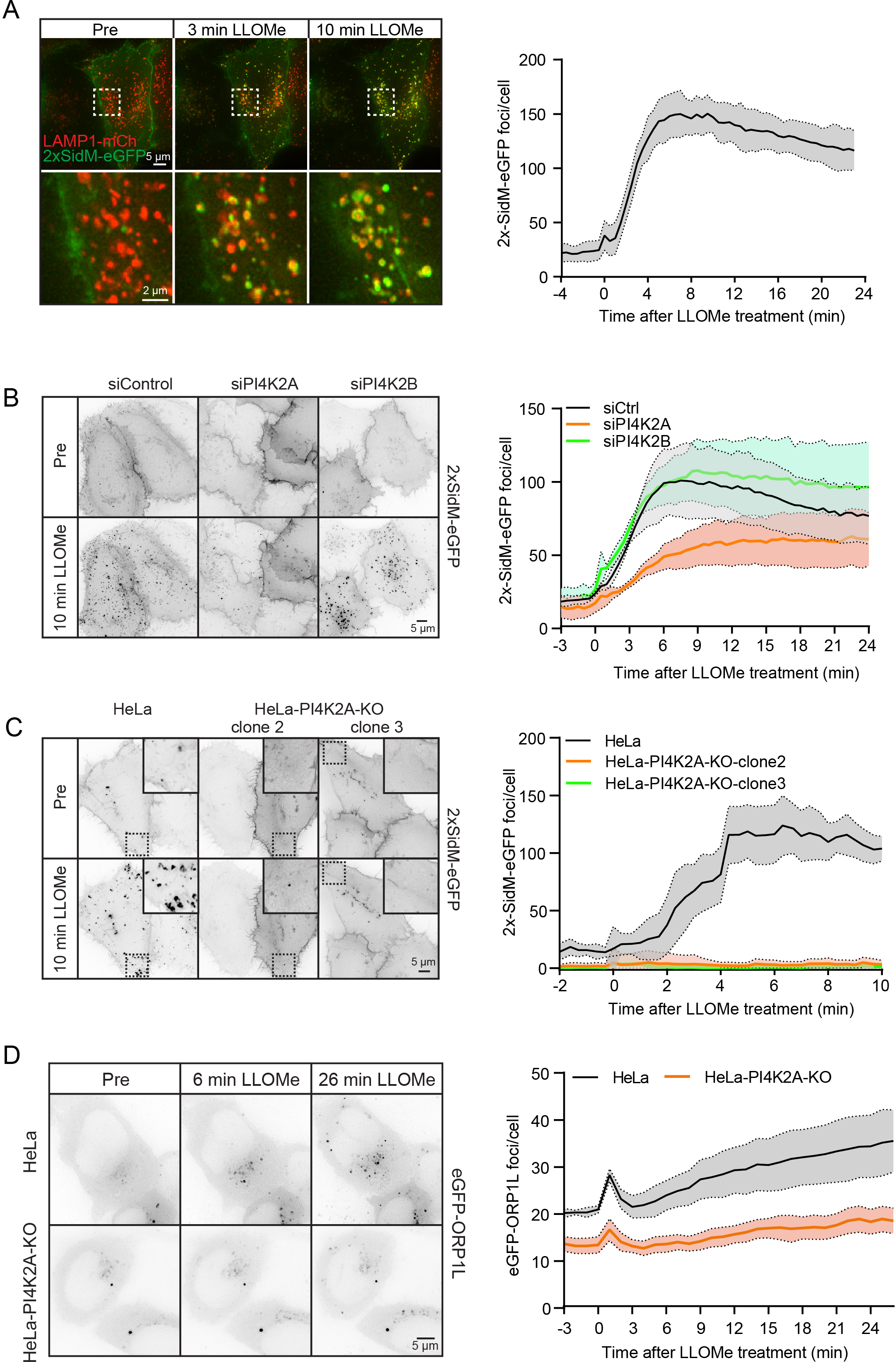
ORP1L is recruited by PI4K2A-dependent accumulation of PtdIns4P on damaged lysosomes. **A** Movie stills from a live-cell imaging experiment showing rapid 2xSidM-eGFP recruitment to damaged lysosomes upon 250 µM LLOMe treatment (indicated in green). Lysosomes were visualized using transiently overexpressed LAMP1-mCherry (indicated in red). Yellow spots show colocalized PtdIns4P probe (2xSidM-eGFP) and LAMP1-mCherry after 3 and 10 min of LLOMe treatment. The graph shows the quantification of 2xSidM-eGFP foci per cell after LLOMe treatment. Error bars denote +/-SEM from n=3 independent live-cell imaging experiments, >20 cells were analyzed per experiment. **B** HeLa cells were transfected with control siRNA or siRNA targeting PI4K2A or PI4K2B for 48 h (KD efficiency is shown in Fig EV6C), and transiently transfected with the 2xSidM-eGFP probe to monitor PtdIns4P recruitment after LLOMe-treatment by live-cell imaging. Representative movie stills show the dynamics of 2xSidM-eGFP recruitment in control, PI4K2A and PI4K2B depleted cells. The graph represents the quantification of 2xSidM-eGFP foci per cell after LLOMe-treatment. Error bars denote +/-SEM from n=3 independent live-cell imaging experiments, >25 cells were analyzed per experiment for each condition. **C** CRISPR-Cas9 mediated knockout of PI4K2A abolishes recruitment of the 2xSidM-eGFP probe (PtdIns4P) to damaged lysosomes. Two different clones (clone 2 and clone 3) were tested. The graph represents the quantification of 2xSidM-eGFP foci per cell after LLOMe-treatment. Error bars denote +/-SD from n=2 independent live-cell imaging experiments, >20 cells were analyzed per experiment for each condition. **D** Representative movie stills from live fluorescence microscopy showing reduced eGFP-ORP1L recruitment in PI4K2A knockout cells compared to wild type HeLa cells upon treatment with 250 µM LLOMe. The graph represents the quantification of eGFP-ORP1L foci per cell after LLOMe treatment. Error bars denote +/-SEM from n=4 independent live-cell imaging experiments, >80 cells were analyzed per experiment for each condition.

PtdIns4P can be produced by 4 different PtdIns 4-kinases (Balla, 2013), and we therefore examined which of these is responsible for the damage-induced formation on lysosomes. For PI4K3A and PI4K3B, inhibitors are commercially available, and we incubated cells expressing 2xSidM-eGFP with GSK-A1 or PI3KIIIB-IN-10, inhibitors of PI4K3A and PI4K3B, respectively. Neither of these inhibitors had any effect on LLOMe-induced PtdIns4P generation, suggesting that PI4K3A and PI4K3B are not involved (Fig EV6A,B). We next examined the possible involvement of PI4K2A and PI4K2B using specific siRNAs to knock down expression of these kinases. Interestingly, whereas there was only a minor effect of siRNA against PI4K2B, siRNA against PI4K2A strongly inhibited LLOMe-induced PtdIns4P accumulation on lysosomes (Figs 3B and EV6C), and CRISPR-Cas9-mediated knockout (KO) of PI4K2A abolished PtdIns4P recruitment as revealed with the 2xSidM probe (Figs 3C and EV6C). Importantly, exogenous expression of PI4K2A-mNG in the PI4K2A knockout cells rescued the recruitment of PtdIns4P after LLOMe treatment (Fig EV6D). This indicates that PI4K2A is specifically activated to produce PtdIns4P on lysosomes in response to damage.

Knowing that PI4K2A produces PtdIns4P on damaged lysosomes, and that the PH domain of ORP1L can bind PtdIns4P (Boutry & Kim, 2021), we investigated whether ORP1L recruitment is affected in PI4K2A knockout cells. Strikingly, live fluorescence microscopy showed that knockout of PI4K2A strongly reduced eGFP-ORP1L recruitment (Fig 3D, Movie EV3). Thus, PI4K2A-mediated formation of PtdIns4P provides a plausible mechanism for recruitment of ORP1L to damaged lysosomes in order to form VAP-containing contact sites with the ER.

### PI4K2A mediates lysosome repair

The conspicuous PI4K2A-mediated accumulation of PtdIns4P and ORP1L on damaged lysosomes begged the question whether PI4K2A is involved in lysosome repair. To address this, we used an assay based on lysosomotropic fluorescent dyes, LysoTracker™, which emit fluorescence when accumulating in acidic lysosomes. Upon damage, lysosome fluorescence ceases because of LysoTracker leakage and pH neutralization, and repair can be monitored with live fluorescence microscopy as recovery of fluorescence (Radulovic *et al*., 2018; Skowyra *et al*., 2018). Using this assay, we found that recovery of LysoTracker fluorescence in lysosomes after LLOMe treatment was strongly inhibited in cells lacking PI4K2A (Fig 4, Movie EV4). This indicates that PI4K2A mediates lysosome repair.

**Figure 4.**
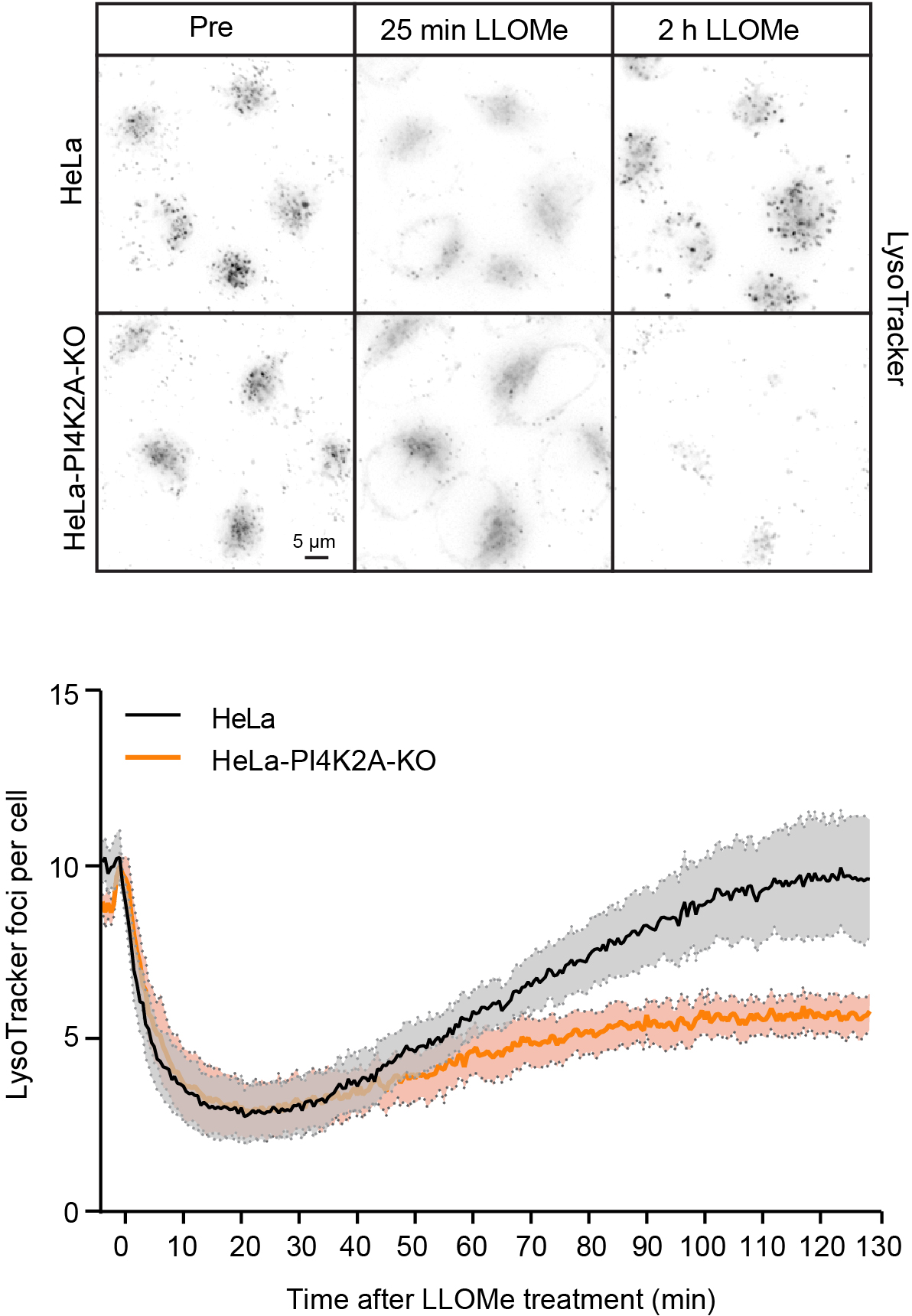
PI4K2A mediates lysosome repair. CRepresentative movie stills from live-cell imaging experiments monitoring Lysotracker recovery after induced lysosomal damage using 250 µM LLOMe. Cells were pre-treated with 75 nM Lysotracker Deep Red for 30 min before addition of LLOMe. The graph represents the quantification of bright Lysotracker positive foci per cell. While the decrease in the number of Lysotracker spots is quickly recovered in HeLa cells, there is a severe impairment in lysosomal repair in HeLa-PI4K2A-KO cells. Error bars denote +/-SEM from n=3 independent experiments, >217 cells were analyzed per experiment for each condition.

### The PI4K2A-mediated lysosome repair pathway is activated independently of ESCRTs

The rapid accumulation of PtdIns4P on lysosomes upon damage occurs with similar kinetics as ESCRT recruitment (Radulovic *et al*., 2018), and hence we asked whether activation of the two repair pathways is connected. Recruitment of the ESCRT-III protein CHMP4B is strongly inhibited by siRNA-mediated knockdown of the upstream ESCRT components ALIX and TSG101 (Radulovic *et al*., 2018), and we therefore monitored whether PtdIns4P accumulation is affected under these conditions. Live microscopy of cells expressing 2xSidM-eGFP showed no effect of ALIX+TSG101 depletion on lysosomal accumulation of the probe (Figs 5A and EV7, Movie EV5), indicating that PI4K2A activation is ESCRT independent.

**Figure 5.**
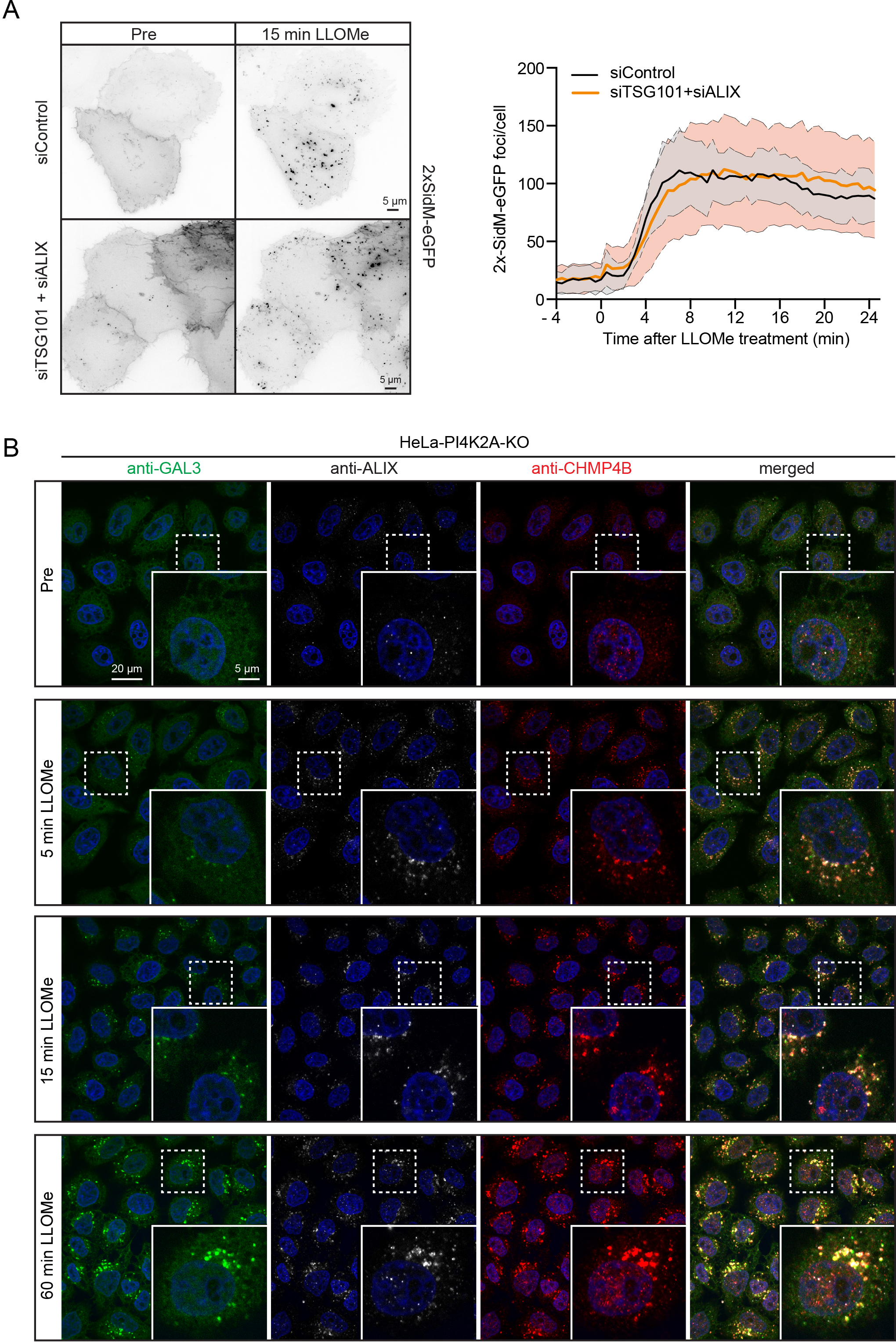
The PI4K2A-mediated lysosome repair pathway is activated independently of ESCRTs. **A** HeLa cells were co-transfected with siRNAs targeting ESCRT proteins TSG101 and ALIX (siTSG101+siALIX) or with non-targeting siRNA control (siControl), and incubated for 24 h before transfection with the PtdIns4P binding probe 2xSidM-eGFP. 24 h post-transfection lysosomes were damaged with 250 µM LLOMe and PtdIns4P recruitment to damaged lysosomes was monitored using the 2xSidM-eGFP probe in live-cell imaging experiments. The graph shows quantification of 2xSidM-eGFP foci per cell in cells co-depleted of TSG101+ALIX and control (siControl) cells. Error bars denote +/-SD from n=2 live-cell imaging experiments, >27 cells were analyzed per experiment for each condition. **B** Representative fluorescence micrographs of HeLa-PI4K2A knockout cells treated with 250 μM LLOMe or equal volume of DMSO (Pre) for 5 min, 15 min, 1 h before fixation and immunostained with Hoechst (blue), anti-CHMP4B (red), anti-GAL3 (green), and anti-ALIX (white).

We next addressed whether ESCRT recruitment depends on PI4K2A. For this purpose we incubated PI4K2A knockout cells with LLOMe, fixed the cells at various time points, and stained with antibodies against the lysosome damage marker GAL3 (Paz *et al*, 2010) as well as ALIX and the ESCRT-III protein CHMP4B (Radulovic *et al*., 2018). Fluorescence microscopy showed that both ALIX and CHMP4B were rapidly recruited, whereas GAL3 recruitment occurred with slower kinetics (Fig 5B), as shown previously (Radulovic *et al*., 2018). This indicates that ESCRT recruitment occurs independently of PI4K2A.

### Cholesterol accumulation on damaged lysosomes depends on PI4K2A, VAPA/B and ORP1L

Since our lipidomics analyses indicated increased cholesterol levels on lysosomes 45 min after damage, we monitored the temporal acquisition of cholesterol on lysosomes. To this end we studied cells expressing an mCherry-tagged cholesterol probe derived from domain 4 (D4) of the bacterial toxin Perfringolysin O (Maekawa, 2017). Interestingly, whereas PtdIns4P accumulated rapidly after LLOMe treatment as revealed with 2xSidM-eGFP, there was a slow but steady accumulation of cholesterol as indicated by the mCherry-D4 probe (Fig 6A, Movie EV6), consistent with the lipidomics data (Fig 1). The LLOMe-induced accumulation of PtdIns4P and cholesterol was abolished in cells depleted for PI4K2A (Fig 6B), indicating that damage-induced cholesterol recruitment requires PI4K2A and PtdIns4P. In VAPA/B knockout cells, PtdIns4P was still recruited, but cholesterol failed to accumulate after damage (Fig 6C). This is consistent with the recruitment of the VAP-dependent cholesterol transporter ORP1L to damaged lysosomes (see Fig 2C), and we therefore investigated whether interference with ORP1L-VAP interactions would affect cholesterol accumulation. For this purpose we studied cells expressing the cholesterol reporter mCherry-D4 and an eGFP-tagged ORP1L mutant incapable of binding to VAP (Vihervaara *et al*, 2011). Live fluorescence microscopy of these cells indicated that the VAP-binding mutation of ORP1L strongly inhibited LLOMe-induced accumulation of cholesterol on lysosomes (Fig 6D, Movie EV7). From these results we conclude that cholesterol accumulation on damaged lysosomes requires both PtdIns4P and the formation of VAP- and ORP1L-containing ER-lysosome contact sites.

**Figure 6.**
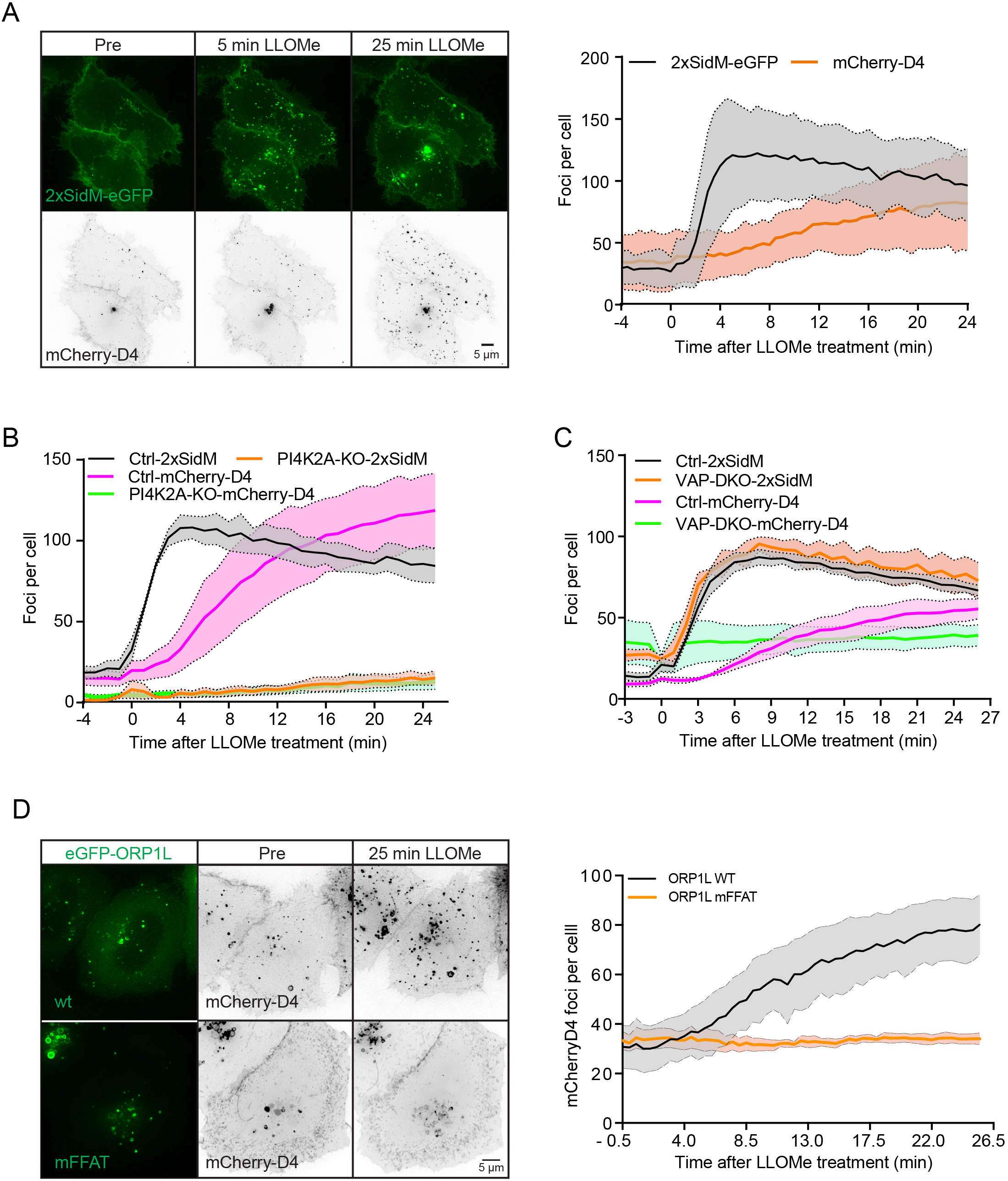
Cholesterol accumulation on damaged lysosomes depends on PI4K2A, VAPA/B and ORP1L. **A** Representative movie stills of a live-cell imaging experiment illustrate faster recruitment of the PtdIns4P probe 2xSidM-eGFP to damaged lysosomes when compared to cholesterol as indicated by the mCherry-D4 probe. The graph shows quantification of 2xSidM-eGFP and mCherry-D4 foci per cell. Error bars denote +/-SD from n=2 live-cell imaging experiments, >26 cells were analyzed per experiment for each condition. **B** Graph showing quantification of mCherry-D4 and 2xSidM-eGFP foci per cell in parental HeLa cells (Ctrl) and PI4K2A knockout cells (PI4K2A-KO). Error bars correspond to +/-SEM from n=3 independent experiments, >30 cells per condition were analyzed. **C** Quantification graph of 2xSidM-eGFP and mCherry-D4 foci per cell in HeLa and VAP double knockout cells. Error bars denote +/-SEM from n=3 independent experiments, >30 cells per condition were analyzed. **D** HeLa cells expressing an eGFP-tagged ORP1L mutant incapable of binding to VAP (mFFAT) show no accumulation of the cholesterol reporter mCherry-D4 upon lysosomal damage induced with 250 µM LLOMe when compared to wildtype (wt) eGFP-ORP1L. The quantification graph shows D4-mCherry foci per cell. Error bars denote +/-SEM from n=4 independent live-cell imaging experiments, >50 cells were analyzed per experiment for each condition.

### Cholesterol protects lysosomes from damage

The correlation between cholesterol accumulation and lysosome damage raised the question whether cholesterol plays a role in the damage response. To investigate this, we took advantage of U18666A, an inhibitor of NPC1, a lysosomal membrane protein that mediates export of cholesterol from lysosomes (Lu *et al*, 2015). U18666A causes accumulation of cholesterol in lysosomes, and we therefore monitored whether such accumulation affects the response to lysosomal membrane damage. Live cell microscopy of HeLa cells stably expressing CHMP4B-eGFP, showed a fast recruitment of CHMP4B to damaged lysosomes in control- and U18666A treated cells, indicating that LLOMe was able to induce an ESCRT-dependent damage response under both conditions (Fig. 7A, Movie EV8). The CHMP4B recruitment was, however, weaker in the U18666A treated cells, and using anti-GAL3 as a damage marker, fluorescence microscopy showed that LLOMe-induced lysosome damage occurred with strongly reduced kinetics in U18666A-treated cells as compared with control cells (Fig 7B,C). This suggests that cholesterol accumulation prevents lysosome damage.

**Figure 7.**
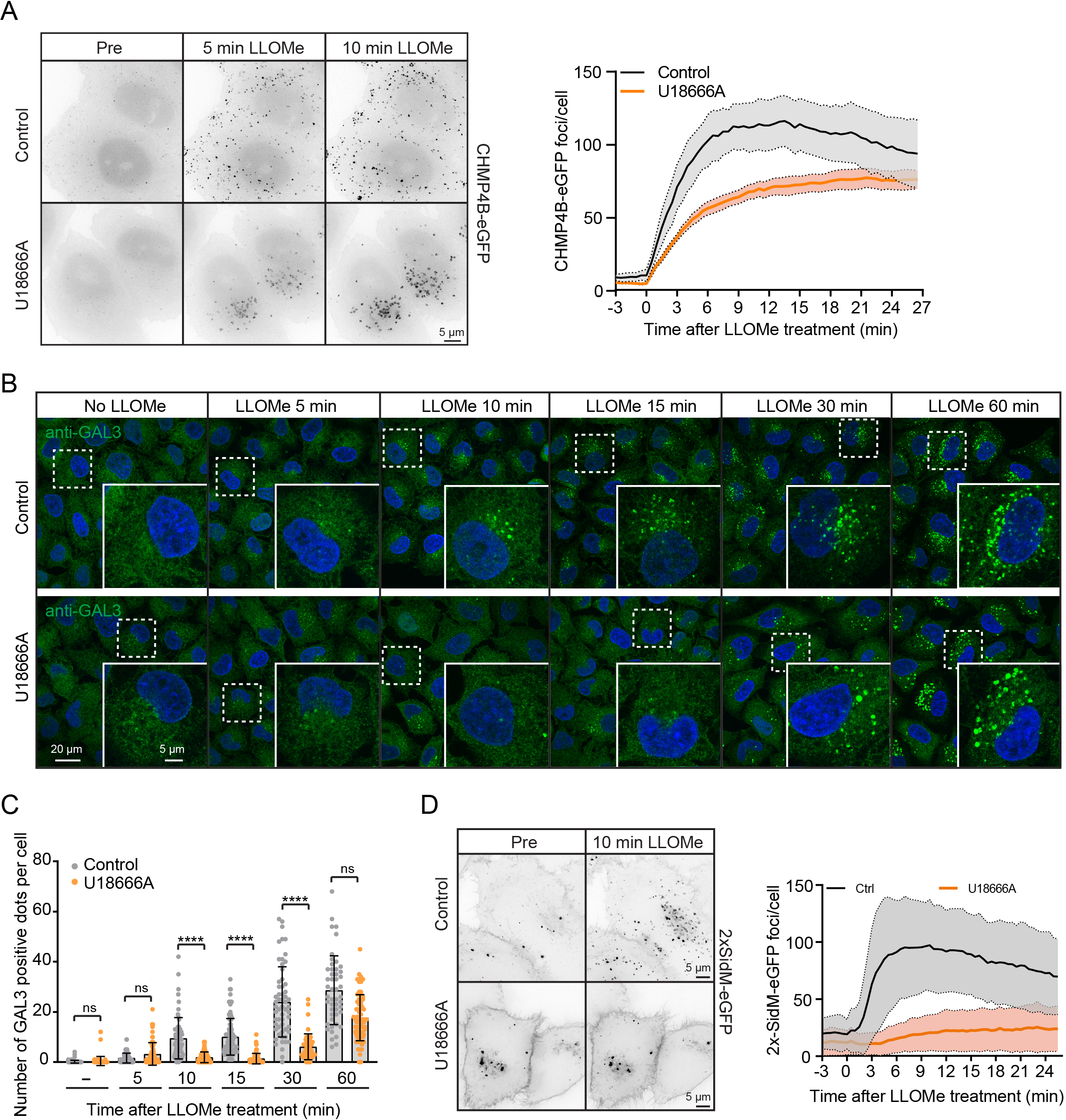
Cholesterol accumulation protects lysosomes from damage. **A** Representative movie stills from live-cell imaging experiments indicate recruitment of CHMP4B-eGFP to damaged lysosomes in control and U18666A treated cells. HeLa cells stably expressing CHMP4B-eGFP were treated with 2 µM U18666A for 17 h before treatment with 250 µM LLOMe. The graph shows quantification of CHMP4B-eGFP foci per cell. Error bars denote +/-SEM from n=4, >30 cells per experiment for each condition. **B** Representative immunofluorescence images illustrate Galectin-3 recruitment kinetics on the damaged lysosomes upon treatment with 2 µM U18666A for 17 h. HeLa cells were treated or not (Control) with U18666A and +/-LLOMe for the indicated time points, fixed and processed for immunofluorescence microscopy. Endogenous levels of Galectin-3 were visualized. **C** Number of Galectin-3 positive dots per cell was quantified from the dataset described in B. Error bars denote +/-SD from n=1, >50 cells per condition. Kruskal-Wallis, Dunn’s post hoc test, **** p<0.0001, ns=not statistically significant. **D** Representative movie stills from live-cell imaging experiments indicate recruitment of the PtdIns4P probe 2xSidM-eGFP to damaged lysosomes in control but not in the U18666A treated cells. HeLa cells were treated with 2 µM U18666A for 17 h before treatment with 250 µM LLOMe. The graph shows quantification of 2xSidM-eGFP foci per cell. Error bars denote +/-SD from n=2, >20 cells per experiment for each condition.

If cholesterol accumulation protects lysosomes from damage, one would expect less recruitment of PtdIns4P to lysosomes in U18666A-treated cells in the presence of LLOMe. This was indeed found to be the case, as U18666A-treated cells showed much weaker increase in lysosomal 2xSidM-eGFP staining after LLOMe incubation than control cells (Fig 7D, Movie EV9). Thus, cholesterol-rich lysosomes seem to be protected from LLOMe-induced damage and are therefore refractory to the PI4K2A-mediated damage response.

### OSBP prevents hyperaccumulation of PtdIns4P on damaged lysosomes, and this promotes cell viability

Since the rapid PtdIns4P recruitment to lysosomes after damage was followed by decreased PtdIns4P levels at later time points (Figs 3A and 6A), there must be cellular mechanisms that remove PtdIns4P from lysosomes. Previous work has shown that the lipid transfer protein OSBP mediates exchange of PtdIns4P for cholesterol at ER-membrane contact sites, followed by dephosphorylation of PtdIns4P into PtdIns by the ER-localized PtdIns4P phosphatase SAC1 (Mesmin *et al*, 2013) and that OSBP ensures a transient pool of PtdIns4P on endosomes (Dong *et al*., 2016). To address whether OSBP might have a similar function at damage-induced ER-lysosome contacts, we first monitored whether eGFP-tagged OSBP is recruited to lysosomes after LLOMe-induced damage. Live fluorescence microscopy showed that this was the case, as eGFP-OSBP gradually accumulated on lysosomes after LLOMe addition (Fig 8A, Movie EV10).

**Figure 8.**
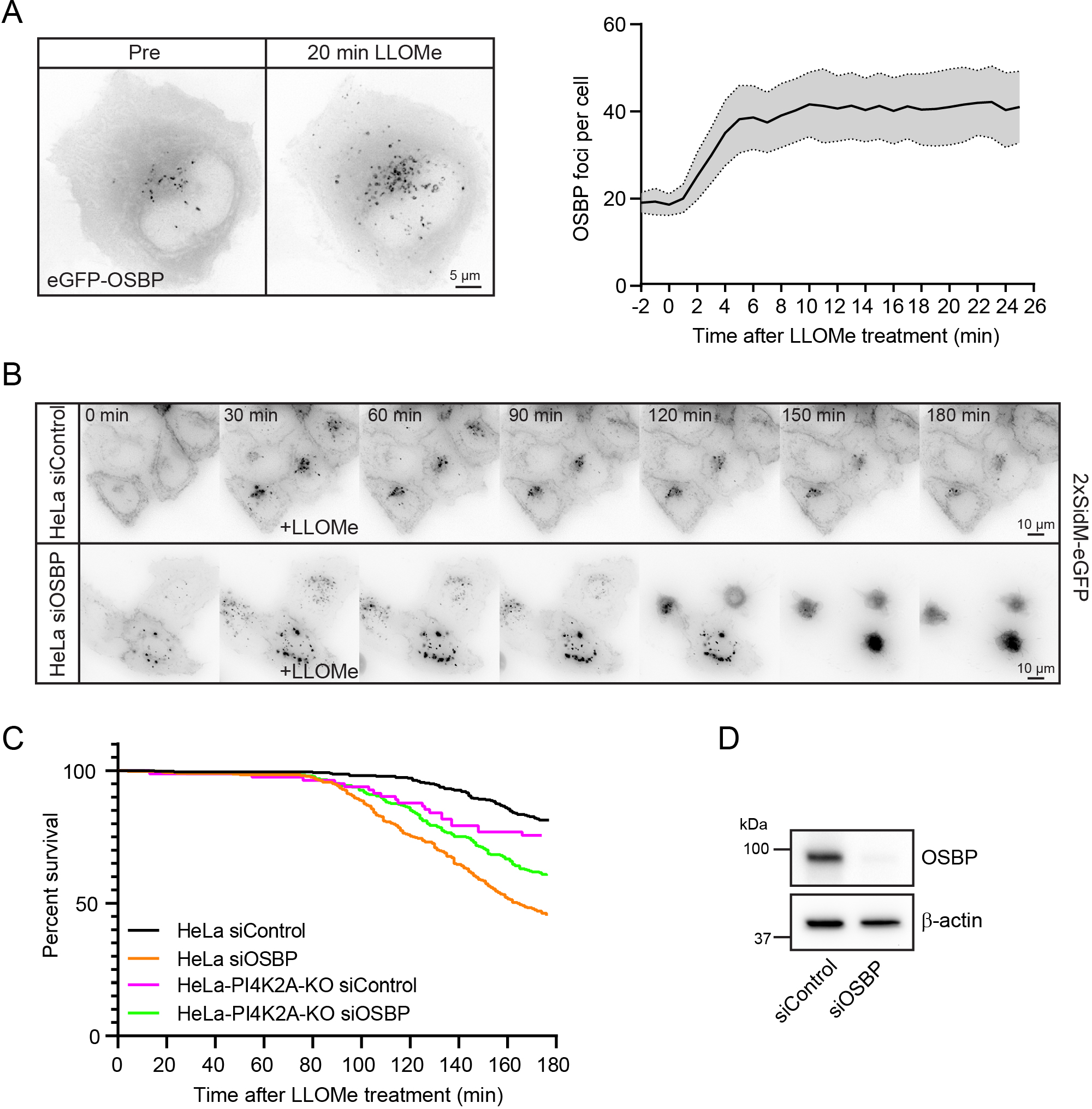
OSBP prevents hyperaccumulation of PtdIns4P on damaged lysosomes, promoting cell viability. **A** Representative movie stills of a live-cell imaging experiment show recruitment dynamics of transiently expressed eGFP-OSBP in HeLa cells treated with 250 µM LLOMe. The graph shows quantifications of OSBP foci per cell. Error bars denote +/-SEM from n=5 independent live-cell imaging experiments, >22 cells per experiment. **B** Representative movie montage from a live-cell fluorescence imaging experiment indicating accumulation of the 2xSidM-eGFP probe in cells depleted of OSBP compared to Control following treatment with 250 µM LLOMe. **C** Kaplan-Meier plot represents decrease in cell viability in siRNA knock down of OSBP (HeLa siOSBP) as compared to control cells (HeLa siControl). The rapid cell death observed in HeLa cells depleted of OSBP was abolished with PtdIns4P depletion (HeLa-PI4K2A-KO siOSBP and HeLa-PI4K2A-KO siControl). The graph represents data from n=3, >100 cells per experiment for HeLa-PI4K2A-KO siOSBP and n=1, >82 cells for HeLa-PI4K2A-KO siControl; n=3, >115 cells per experiment for HeLa siControl and n=3, >120 cells per experiment for HeLa siOSBP. **D** Knock down efficiency of siRNA depletion of OSBP in HeLa cells as detected by Western blot. β-actin used as a loading control.

In order to investigate whether OSBP is involved in PtdIns4P removal from damaged lysosomes, we used siRNA against OSBP to knock down its expression and used 2xSidM-eGFP and live fluorescence microscopy to monitor PtdIns4P levels on lysosomes after LLOMe-induced damage. Depletion of OSBP did in fact cause a strong accumulation of 2xSidM-eGFP on damaged lysosomes, indicating that OSBP plays a role in removal of PtdIns4P. Surprisingly, accumulation of PtdIns4P in response to OSBP depletion was accompanied by cell death as revealed by rounding and detachment of the OSBP-depleted cells at about 120 minutes after LLOMe addition (Fig 8B-D, Movie EV11).

To study if cell death was caused by accumulation of PtdIns4P, we monitored whether knockout of PI4K2A prevented the rapid cell death observed upon OSBP depletion. Indeed, although PI4K2A KO cells depleted for OSBP eventually died after LLOMe addition, as expected after lysosome damage, the rapid death observed with PtdIns4P depletion was abolished (Fig 8C). These results indicate that OSBP-mediated removal of PtdIns4P after PtdIns4P-mediated membrane repair is essential for cell viability.

## DISCUSSION

A number of recent publications have demonstrated that cells are equipped with alternative mechanisms to deal with damaged lysosomes (Herbst *et al*, 2020; Hung *et al*., 2013; Jia *et al*, 2020a; Jia *et al*, 2020b; Lopez-Jimenez *et al*., 2018; Maejima *et al*., 2013; Niekamp *et al*., 2022; Radulovic *et al*., 2018; Skowyra *et al*., 2018). We have uncovered a new mechanism that initiates with PI4K2A-mediated generation of PtdIns4P on the membrane of the damaged lysosome. This is followed by recruitment of ORP1L and VAP-containing contacts between the damaged lysosomes and the ER. ORP1L transfers cholesterol to the lysosome membrane to augment its repair (Fig 9).

**Figure 9.**
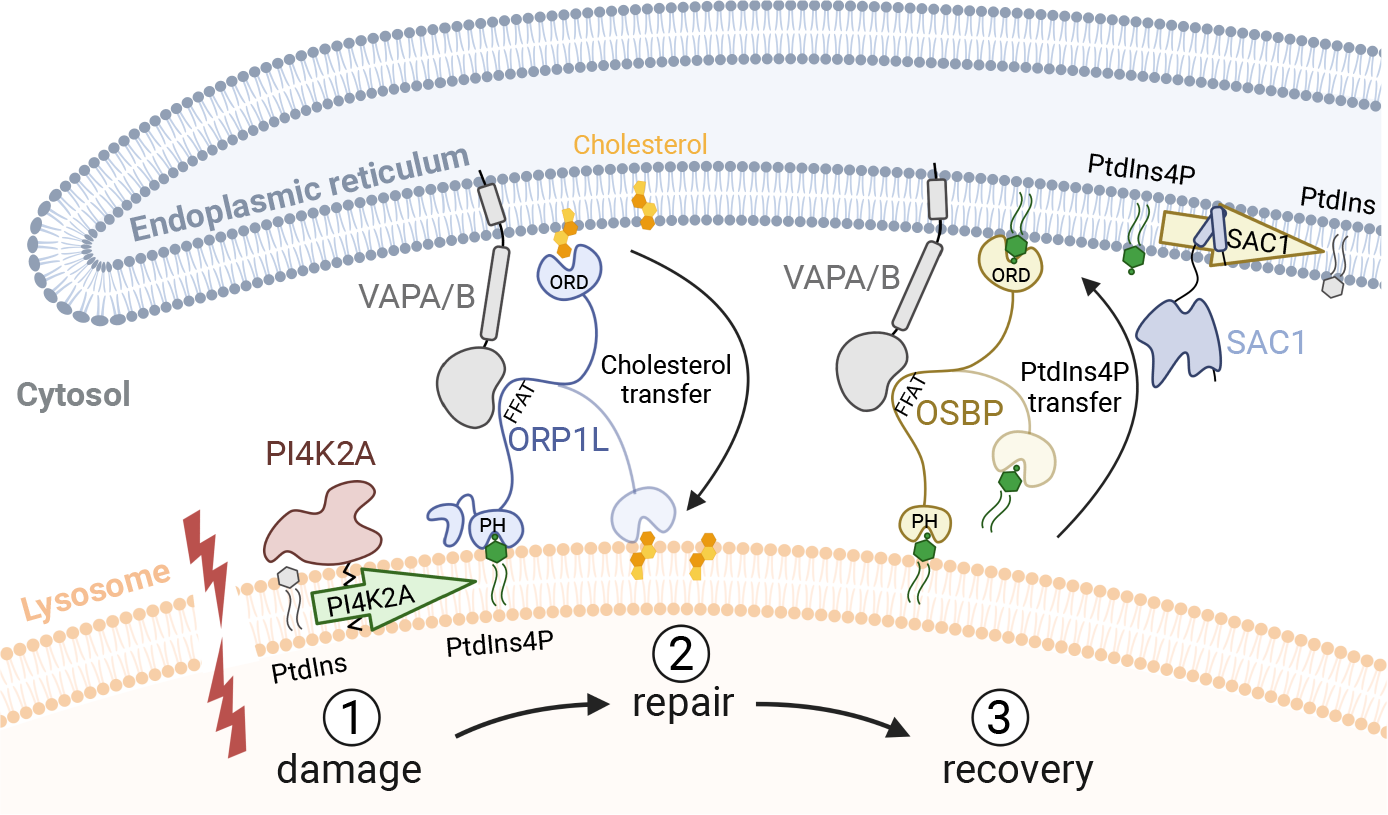
Model for ER-mediated lysosome repair. Damage (1) causes activation of PI4K2A to produce PtdIns4P on the lysosome membrane. This causes formation of contact sites via ORP1L, OSBP and other PtdIns4P-binding ORP proteins that bind to VAPA/B in the ER membrane via their FFAT motives and to PtdIns4P on the lysosome via their PH domains. Transfer of cholesterol from the ER, mediated by ORP1L, promotes membrane repair (2). Excess PtdIns4P is transported from the lysosome to the ER by OSBP (3) and thereafter dephosphorylated by SAC1 (the latter inferred from (Mesmin *et al*., 2013)). The lipid binding ORD domains of ORP1L and OSBP are indicated.

While this manuscript was in preparation, another paper reporting the involvement of PI4K2A, PtdIns4P and ER contacts in lysosome repair was published (Tan & Finkel, 2022). This paper identified three PS transfer proteins, ORP9, ORP10 and ORP11, as being recruited to damaged lysosomes by PtdIns4P, and also made the interesting observation that the lipid channel protein ATG2 is activated by PS for repair of damaged lysosomes. The involvement of ORP9, ORP10, ORP11 and ATG2 provides a plausible explanation for the increased PS levels on damaged lysosomes that we detected in our lipidomics analyses. Here we have provided evidence that ORP1L-mediated cholesterol transfer is also important for repair.

The striking increase in ER-lysosome contact sites observed by electron microscopy shortly after LLOMe addition indicates that contact sites are rapidly formed upon lysosome damage. For this reason, we cannot formally exclude the possibility that detected changes in the lysosomal lipidome upon LLOMe might be in part due to contamination of the immunoisolated lysosomes with ER membranes. Indeed, the increase in PC levels detected at 10 minutes after LLOMe might be due to ER contamination since PC is a major lipid in ER membranes (Scrima *et al*., 2022), even though Western blotting with ER antibodies revealed minimal contamination. However, the increase in PS and cholesterol levels at 45 minutes after lysosome damage cannot be explained by ER contamination, and the increase in cholesterol was verified independently with a cholesterol-binding probe and correlates well with the acquisition of the cholesterol transporter ORP1L detected by live microscopy.

What is the role of PtdIns4P in lysosome repair? The most obvious function would be to recruit PtdIns4P-binding proteins like ORP9, ORP10, ORP11 and ORP1L to establish contacts with the ER via the VAP-binding domains of the ORP proteins. However, elegant work on ER-Golgi contacts has revealed an additional function for PtdIns4P in membrane contact sites, namely in driving cholesterol transfer. In particular, OSBP has been shown to transfer cholesterol from ER to Golgi membranes in exchange for PtdIns4P, and dephosphorylation of PtdIns4P in the ER membrane serves to maintain a concentration gradient that drives the exchange (Mesmin *et al*., 2013). We found that OSBP depletion causes a strong accumulation of PtdIns4P on damaged lysosomes, suggesting that OSBP performs a similar function in ER-lysosome contact sites. Therefore, OSBP not only mediates cholesterol transfer as proposed (Tan & Finkel, 2022), but also serves to balance lysosomal PtdIns4P levels.

Our observation that lysosomal PtdIns4P accumulation in the absence of OSBP was accompanied by cell death illustrates the importance of PtdIns4P removal, although we still need to define why PtdIns4P accumulation is so hazardous to the cells suffering from lysosomal injury. PtdIns4P accumulation on lysosomes would induce increased and prolonged ER-endosome contact sites, since many contact site proteins depend on PtdIns4P, and this might affect lysosome function. In addition, accumulation of endolysosomal PtdIns4P causes accumulation of membrane-associated actin leading to impaired recycling from endolysosomal tubules (Dong *et al*., 2016).

It also needs to be explained exactly how transfer of PS and cholesterol from the ER would mediate lysosome repair, and a relevant question is how the ER-mediated repair relates to the previously identified ESCRT- and ceramide-dependent repair mechanisms (Niekamp *et al*., 2022; Radulovic *et al*., 2018; Skowyra *et al*., 2018). The latter two mechanisms appear to be independent (Niekamp *et al*., 2022), and both we and others (Tan & Finkel, 2022) have found that activation of the ESCRT- and ER-mediated repair occurs independently of each other. However, independent activation mechanisms do not exclude the possibility that the alternative repair mechanisms cross-talk at some level. For instance, if ESCRTs and ceramide mediate inward budding of damaged membrane areas to be degraded in the lysosome lumen (Niekamp *et al*., 2022; Radulovic *et al*., 2018), addition of more lipid would be required to maintain the area of the lysosome membrane, and ER-mediated lipid transfer might play a role here. Further studies are needed in order to clarify these issues.

## ACKNOWLEDGEMENTS

The Core Facilities for Advanced Light Microscopy and Advanced Electron Microscopy at Oslo University Hospital are acknowledged for providing access to and training on relevant microscopes. We thank Ulrikke Dahl Brinch and Catherine Sem Wegner for excellent technical help with sample preparation for electron microscopy. We thank Pietro De Camilli for kindly providing VAPA/B knockout cells and Matthew Yoke Wui Ng and Kia Wee Tan for providing plasmid constructs. H.S. was supported by the Norwegian Cancer Society (project number 182698), the South-Eastern Norway Regional Health Authority (project number 2016087), the Research Council of Norway (project number 302994) and the European Research Council (Advanced Grant number 788954). M.J. was supported by grants from the Danish National Research Foundation (DNRF125) and the Novo Nordisk Foundation (NNF17OC0029432), and K.M. was supported by the Independent Research Fund Denmark (6108–00542B). This work was partly supported by the Research Council of Norway through its Centres of Excellence funding scheme (project number 262652). Figures were created using Adobe Illustrator CS6 and BioRender (https://biorender.com/).

## MATERIALS AND METHODS

### Reagents, cell culture, and stable cell lines

The lysosomotropic drug used in this study, L-leucyl-L-leucine methyl ester (cat. no. 16008) was purchased from Cayman Chemical and used at 250 μM. LLOMe was dissolved in dimethyl sulfoxide (DMSO) and stored at −20°C. Other reagents used in this study: LysoTracker Deep Red (cat. no. 12492) for live-cell imaging was purchased from Molecular Probes and was used at 75 nM, GSK-A1 (cat. no. SML2453-5MG) was purchased from Sigma-Aldrich and was used at 10 nM, PI4KIIIB-IN-10 (cat. no. HY-100198) was purchased from MedChemExpress and was used at 25 nM, U18666A (cat. no. U3633-5MG) was purchased from Merck Life Science (Sigma) and used at 2 µM.

Human HeLa “Kyoto” cells were maintained in DMEM (Gibco) medium, supplemented with 10% fetal bovine serum (FBS), 5 U/ml penicillin, and 50 μg/ml streptomycin. Cells were maintained at 37°C supplemented with 5% CO_2_. A stable HeLa cell line expressing CHMP4B-eGFP was obtained from Anthony A. Hyman (Max Planck Institute for Molecular Cell Biology and Genetics, Dresden, Germany). The HeLa-VAP-DKO stable cell line was obtained from Pietro De Camilli (Yale School of Medicine, New Haven, USA).

### CRISPR/Cas9-mediated deletion of PI4K2A

CRISPR/Cas9-mediated deletion of PI4K2A in HeLa ‘‘Kyoto’’ cells was introduced using CRISPR/Cas9. Guide RNAs were designed using Benchling software (www.benchling.com). For deletion of PI4K2A the following guide was used gRNA: 5’-TTAATCCTAAGTGGACCAAG-3’. pX459-derived plasmids encoding both Cas9 and the gRNA were transfected using Fugene 6 (cat. no. E2692 from Promega). 24 h post-transfection puromycin was used to select for transfected cells, these cells were further sorted to obtain single-cell derived colonies, which were expanded and further characterized. Clones lacking PI4K2A were identified through Western blotting.

### siRNA transfections

Silencer Select siRNAs against TSG101 (#1, 5′-CCGUUUAGAUCAAGAAGUA-3′; #2, 5′-CCUCCAGUCUUCUCUCGUC-3′), ALIX (#1, 5′-GCAGUGAGGUUGUAAAUGU-3′; #2, 5′-CCUGGAUAAUGAUGAAGGA-3′), OSBP (predesigned, cat. no. 4392420) and nontargeting control siRNA (predesigned, cat. no. 4390844) were purchased from Ambion. ON-TARGET plus Human PI4K2A siRNA (predesigned, cat. no. #1 J-006770-06-0002, #2 J-006770-07-0002) and ON-TARGET plus Human PI4K2B siRNA (predesigned, cat. no. #1 J-006769-06-0002, #2 J-006769-07-0002) were purchased from Dharmacon. Cells at 50% confluency were transfected with 20–40 nM final siRNA concentration using Lipofectamine RNAiMax transfection reagent (Life Technologies) according to the manufacturer’s instructions and harvested after 48 h.

### Plasmid constructs

mCherry-P4M-SidM was a gift from Tamas Balla (Addgene plasmid # 51471, (Hammond *et al*., 2014)), GFP-P4M-SidMx2 was a gift from Tamas Balla (Addgene plasmid # 51472 (Hammond *et al*., 2014)), GFP-C1-PLCdelta-PH was a gift from Tobias Meyer (Addgene plasmid # 21179,(Stauffer *et al*, 1998)), mCherry-D4 was a gift from Matthew Ng. PI4K2A-mNG was a gift from Kia Wee Tan. pLJM1-FLAG-GFP-OSBP was a gift from Roberto Zoncu (Addgene plasmid # 134659 (Lim *et al*, 2019)), eGFP-ORP1L was described previously (Johansson *et al*, 2007).

### Immunofluorescence staining and antibodies

Cells seeded on coverslips were fixed in 4% EM-grade formaldehyde in phosphate-buffered saline (PBS) for 10 min. Cells were washed twice in PBS and once in PBS containing 0.05% saponin to permeabilize the cells before staining with the indicated primary antibodies for 1 h. Prior to staining with the secondary antibodies for 1 h, cells were washed three times in PBS containing 0.05% saponin. Cells were mounted in Mowiol containing 2 mg/ml Hoechst 33342 (Sigma-Aldrich). Rabbit anti-ALIX and rabbit anti-CHMP4B antibodies were described previously (Christ *et al*, 2016). Human Galectin-3 Alexa Fluor 488-conjugated antibody (cat. no. IC1154G) was purchased from R&D Systems. Mouse anti-LAMP1 (cat. no. H4A3) was purchased from Developmental Studies Hybridoma Bank, mouse anti-β-actin (cat. no. A5316) from Sigma-Aldrich, and mouse anti-TSG101 (cat. no. 612697) from BD Transduction Laboratories. Rabbit anti-Calnexin antibody (cat. no. 2679S) was purchased from Cell Signaling Technology, mouse anti-TOM20 antibody (cat. no. 612278) was purchased from BD Biosciences. All secondary antibodies used for immunofluorescence studies and Western blotting were obtained from Jacksons ImmunoResearch Laboratories or from Molecular Probes (Life Technologies).

### Western blotting

Cells were washed with cold PBS and lysed in 2× sample buffer (125 mM Tris–HCl, pH 6.8, 4% SDS, 20% glycerol, 200 mM DTT, and 0.004% bromophenol blue). Whole-cell lysates were subjected to SDS–PAGE on 4–20% gradient gels (Mini-PROTEAN TGX; Bio-Rad). Proteins were transferred to polyvinylidene difluoride (PVDF) membranes (Trans-Blot^®^ Turbo™ LF PVDF, Bio-Rad) followed by blocking in 5% fat-free milk powder and antibody incubation in 5% fat-free milk powder in Tris-buffered saline with 0.1% Tween-20. Membranes incubated with horseradish peroxidase-conjugated antibodies were developed using Clarity Western ECL Substrate Solutions (Bio-Rad) with a ChemiDoc XRS+ imaging system (Bio-Rad).

### Confocal fluorescence microscopy

Stained coverslips were examined with a Zeiss LSM 780 confocal microscope (Carl Zeiss) equipped with an Ar laser multiline (458/488/514 nm), a DPSS-561 10 (561 nm), a laser diode 405-30 CW (405 nm), and a HeNe laser (633 nm). The objective used was a Zeiss Plan-Apochromat 63×/1.40 Oil DIC M27 (Carl Zeiss). Intensity settings for the relevant channels were kept constant during imaging. Image processing was performed with ImageJ software (National Institutes of Health).

### Live-cell imaging

Cells seeded in MatTek 35-mm petri dish, 20-mm Microwell No. 1.5 coverglass were imaged on a DeltaVision microscope (Applied Precision) equipped with Elite TruLight Illumination System, a CoolSNAP HQ2 camera, and a 60× Plan-Apochromat (1.42 NA) lens. For temperature control during live observation, the microscope stage was kept at 37°C by a temperature-controlled incubation chamber. Time-lapse images (6 *z*-sections 2.2 μm apart) were acquired every 1–3 min over a total time period of 4–6 h and deconvolved using the softWoRx software (Applied Precision). In addition, DeltaVision OMX V4 microscope equipped with three PCO.edge sCMOS cameras, a solid-state light source, a 60× 1.42 NA objective, and a laser-based autofocus was used. Environmental control was provided by a heated stage and an objective heater (20-20 Technologies). Images were deconvolved using softWoRx software and processed in ImageJ/FIJI.

### Image analysis and post-processing

Images were analyzed in Fiji/ImageJ using custom python scripts. Briefly, images were filtered to remove noise and foci of different proteins were segmented by manual thresholding and objects were then identified and scored by the “Analyze Particles” function of ImageJ. Manual thresholds were chosen to detect prominent LLOMe-induced structures; therefore, only spots above a certain intensity were scored. For live-cell movies, foci counts were normalized to the cell number. For fixed cell analysis, nuclei were segmented and counted, and all foci counts were normalized to the number of nuclei within a given field of view.

### Transmission electron microscopy (TEM)

Cells were grown on glass-coverslips (100.000 cells per well in 12-well plates) and incubated with BSA-gold for 16 hours to identify lysosomes. They were fixed at different time-points after LLOMe treatment, with 2% glutaraldehyde in 0.1 M PHEM buffer. Fixative was removed, coverslips were washed and postfixation was done in 1% Osmium (EMS, 19134) /1.5% Ferricyanide/0.1 M PHEM for 1 h. Cells were thoroughly washed in water to remove Osmium residues. The cells were contrasted using 4% uranyl acetate (SPI, 02624-AB) and dehydrated in a graded ethanol series, embedded in Epon and polymerized for 48 h. Ultrathin sections of 100 nm were cut using an Ultracut UCT ultramicrotome (Leica, Austria) and transferred onto formvar/carbon coated grids. All prepared sections were observed at 80 kV in a JEOL-JEM 1230 electron microscope and images were recorded using iTEM software with a Morada camera (Olympus, Münster, Germany).

### Lysosome purification and lipidomics

Lysosomes were purified and their lipid contents were analyzed with quantitative mass spectrometry-based shotgun lipidomics as described previously (Nielsen *et al*., 2020; Stahl-Meyer *et al*., 2022). In brief, the post-nuclear fractions prepared from the LLOMe-treated or untreated cells were incubated with anti-LAMP-1 primary antibody (Abcam, #ab24170) or IgG control (Invitrogen, Waltham, MA, USA, #31235) and then with anti-Rabbit IgG MicroBeads (Miltenyi Biotec, #130-048-602). The post-nuclear fractions were loaded on magnetic columns (Miltenyi Biotec, Bergisch Gladbach, Germany, #130-042-401) to capture and elute lysosomes associated with MicroBeads. Aliquots of the eluates and the corresponding whole cell lysates were centrifuged at 21,100× *g* for 20 min, and the pellets were resuspended in 155 mM NH_4_HCO_3_ for shotgun lipidomics.

For shotgun lipidomics, a series of lipid internal standards in appropriate and known quantities (Stahl-Meyer *et al*., 2022) were added to each sample prior to lipid extraction using a modified Bligh and Dyer protocol. The lipid extracts were mixed with positive (13.3 mM ammonium bicarbonate in 2-propanol) or negative (0.2% (*v*/*v*) tri-ethyl-amine in chloroform:methanol 1:5 (*v*/*v*)) ionization solvents. The lipidomics analysis was performed on a quadrupole-Orbitrap mass spectrometer Q Exactive equipped with a TriVersa NanoMate to allow direct infusion and ionization of the samples. The data were acquired through cycles of MS and MS/MS scans in both positive and negative ion modes. The acquired ion spectra were processed using a Python-based software, LipidXplorer (Herzog *et al*, 2012), which reported the lipid species identified based on sets of criteria defined for the individual lipid classes as well as their *m/z* values and the intensities of associated precursor and fragment ions (Stahl-Meyer *et al*., 2022). A subsequent sorting and quantification of lipid species were achieved using an in-house built R-based suite of scripts named LipidQ (https://github.com/ELELAB/lipidQ) and manual evaluation of the spectra. The absolute molar quantities were calculated based on intensity values of sample-derived lipids and their respective internal lipid standards.

Species of phosphatidylinositol phosphate (PIP) were identified in the negative ion mode using LipidXplorer based on the following criteria; MS1 detection (tolerance 6 ppm) of [M-2H]^2-^ precursor ion and MS2 detection (tolerance 0.0035 Da) of dehydrated phosphorylinositol headgroup, dehydrated glycerol phosphate, and a pair of carboxylate anions of fatty acids matching the sum composition of the associated precursor ion. Intensities of the precursor ions and the fragment ions were retrieved using LipidXplorer.

### Lipidomics Data Analysis and Statistics

The molar quantity of lipids were expressed relative to the sum of the molar quantities of all identified lipid species in the sample (mol%) unless otherwise specified. Lipid species were only included in the further analyses if the determined lipid species were detected in at least 75% of the replicates. Heatmaps were produced using the “pheatmap” package in R (version 4.0.3.) (Nielsen *et al*., 2020). For statistical analyses of lipid levels, *t*-tests with Benjamin-Hochberg (BH) correction for multiple testing were performed in R and adjusted *p*-values (*q*-values) were determined. Lipid were considered to be at statistically significantly different levels between two sample groups if the adjusted *p*-values were below 0.05. Lipid nomenclatures were as previously used (Nielsen *et al*., 2020).

### Statistical analysis

The number of individual experiments and the number of cells analyzed are indicated in the figure legends. We tested our datasets for normal distribution and chose an appropriate test accordingly using GraphPad Prism version 5.01. The statistical tests performed for each experiment are specified in the figure legends. *P*-values are indicated for each experiment, and *P*-values below 0.05 were regarded as significant. In instances of multiple means comparisons, we used one-way analysis of variance (ANOVA) followed by the Dunnett’s post hoc test to determine statistical significance.

## EXPANDED VIEW FIGURE LEGENDS

**Figure EV1.**
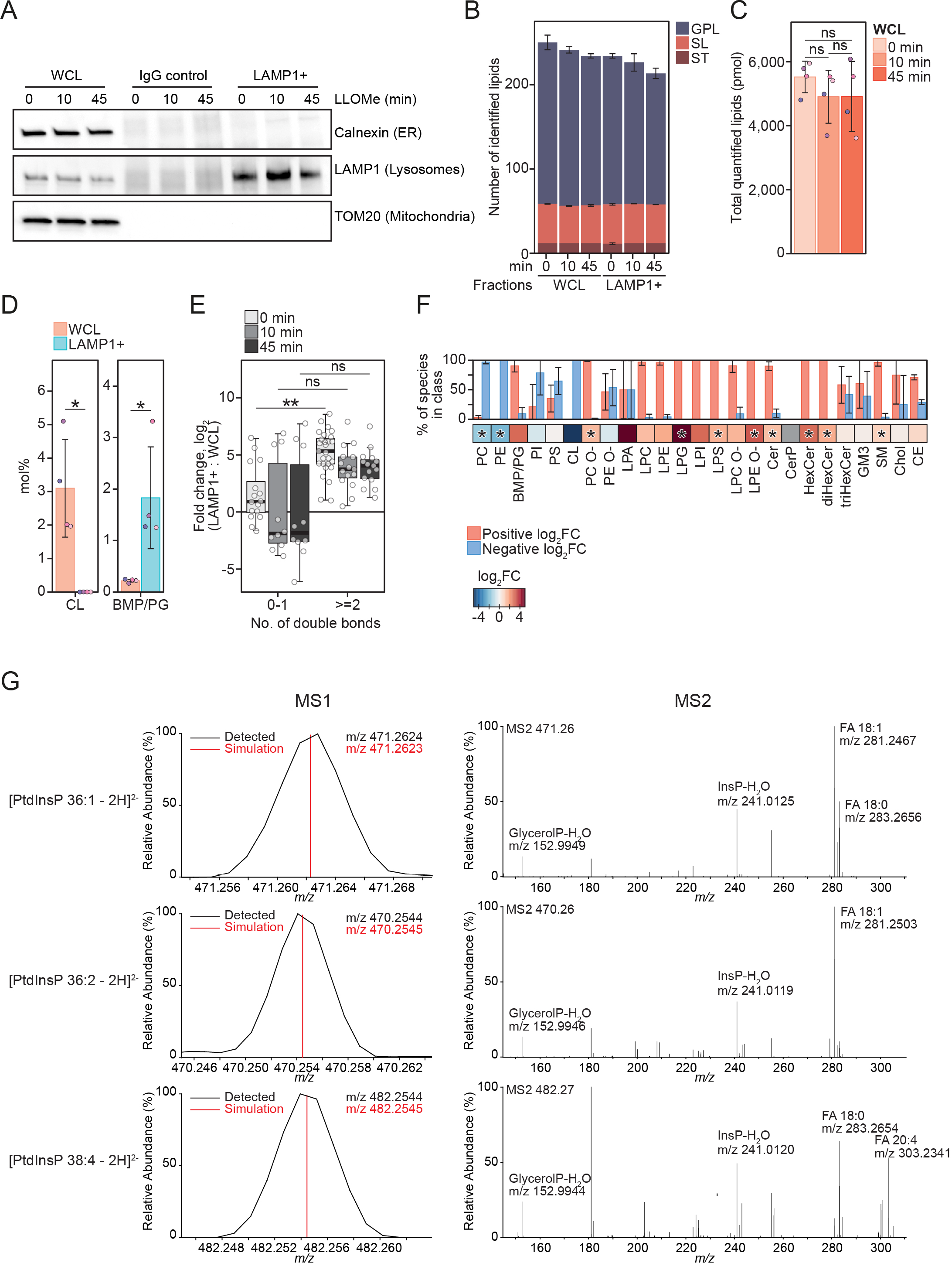
**A** Representative Western blot showing purity of immuno-affinity purified lysosomal (LAMP1+) and negative control (IgG Control) treated with 250 µM LLOMe for 0, 10 min and 45 min. Membrane was probed using anti-TOM20 and anti-Calnexin antibodies to verify the absence of mitochondrial and ER membrane, respectively. Enrichment of lysosomes was detected using anti-LAMP1 antibody. **B** Numbers of lipid species identified in the produced WCLs and LAMP1+ fraction, belonging to the lipid categories glycerophospholipids (GPL), sphingolipids (SL), and sterol lipids (ST). **C** Total molar quantities of lipids identified in aliquots of the whole cell lysates (WCL) produced during lysosome purification. **D** Mol% values of the cardiolipin (CL) and bis(monoacylglycero)phosphate/phosphatidylglycerol (BMP/PG) classes in the LAMP1+ fraction and WCLs of untreated HeLa cells. BMP and PG classes are isobaric and reported together as BMP/PG. **E** Enrichment of individual BMP/PG species after purification (log_2_-transformed fold change, mol% in LAMP1+: mol% in WCL), presented as box plots with individual species as circles. Species are grouped according to the total number of acyl double bonds. **F** Enrichment of monitored lipid classes (log_2_-transformed fold change, mol% in LAMP1+: mol% in WCL) after purification of lysosomes from untreated cells, presented in a heatmap. The bar plot depicts percentage of species in the individual classes having positive or negative values of log_2_ fold changes. **G** Representative mass spectra of three PtdInsP species. Precursor ions detected in the MS1 and their theoretical *m/z* values (MS1) and MS2 spectra obtained after their fragmentations, displaying the relative intensities of the ions. Multiple t-tests were performed to assess changes between means of pmol (C) and mol% (D and F). Resultant *p*-values were corrected for multiple testing using Benjamini-Hochberg correction with an adjusted *p*-value (q-value) of 0.05. * q < 0.05. N=3-4. *Abbreviations*: FA, fatty acid; glycerolP, glycerol phosphate; InsP, phosphorylinositol headgroup. Additional abbreviations are as in Fig 1.

**Figure EV2.**
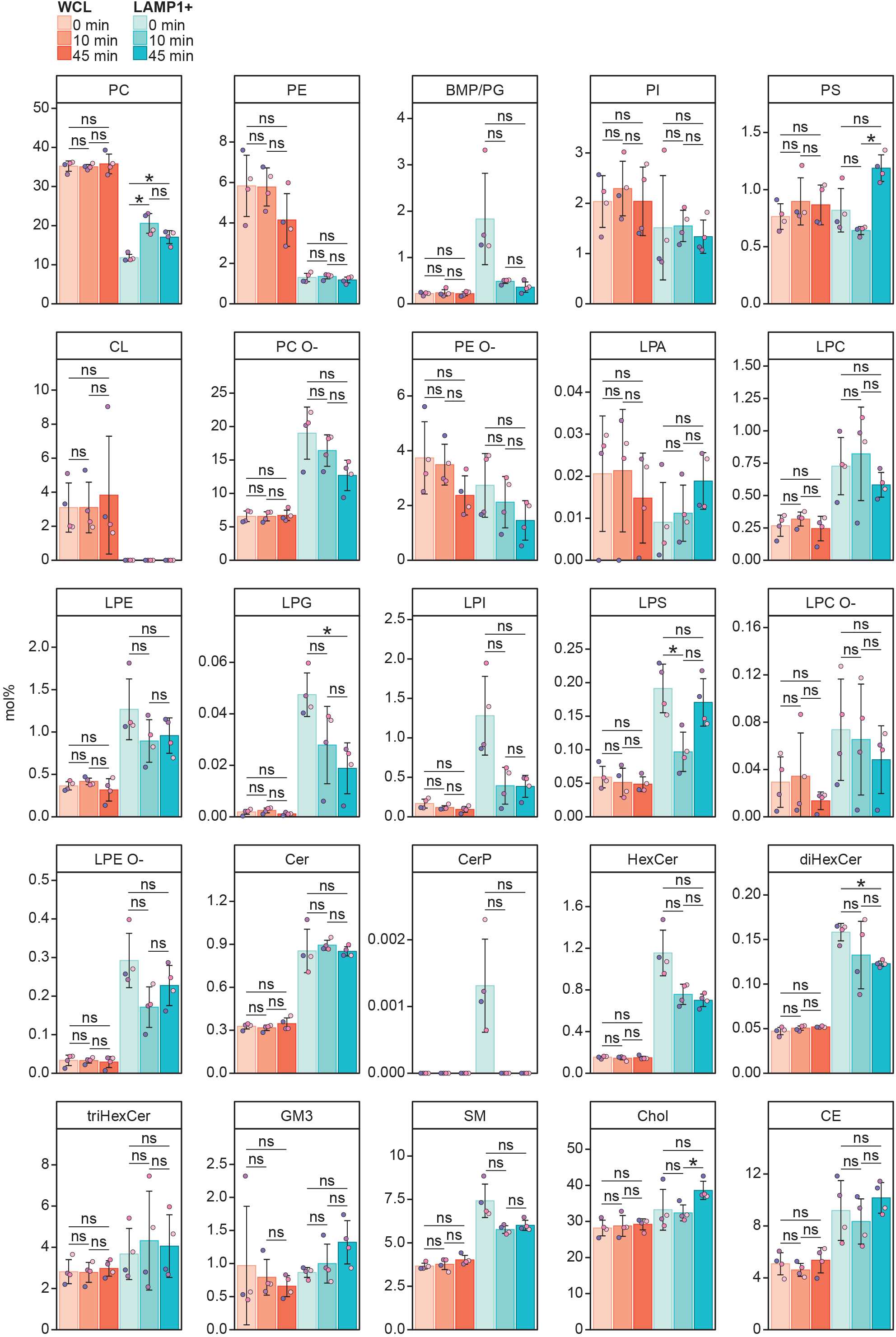
Mol% of all monitored lipid classes in the WCLs and LAMP1+ fractions from cells untreated or treated with LLOMe for 10 or 45 min. Multiple t-tests were performed to assess changes between means. Resultant *p*-values were corrected for multiple testing using Benjamini-Hochberg correction and differences were considered statistically significantly changed with adjusted *p*-values (q-values) of 0.05 or less. * q < 0.05. N=3-4. Abbreviations are as in Fig 1.

**Figure EV3.**
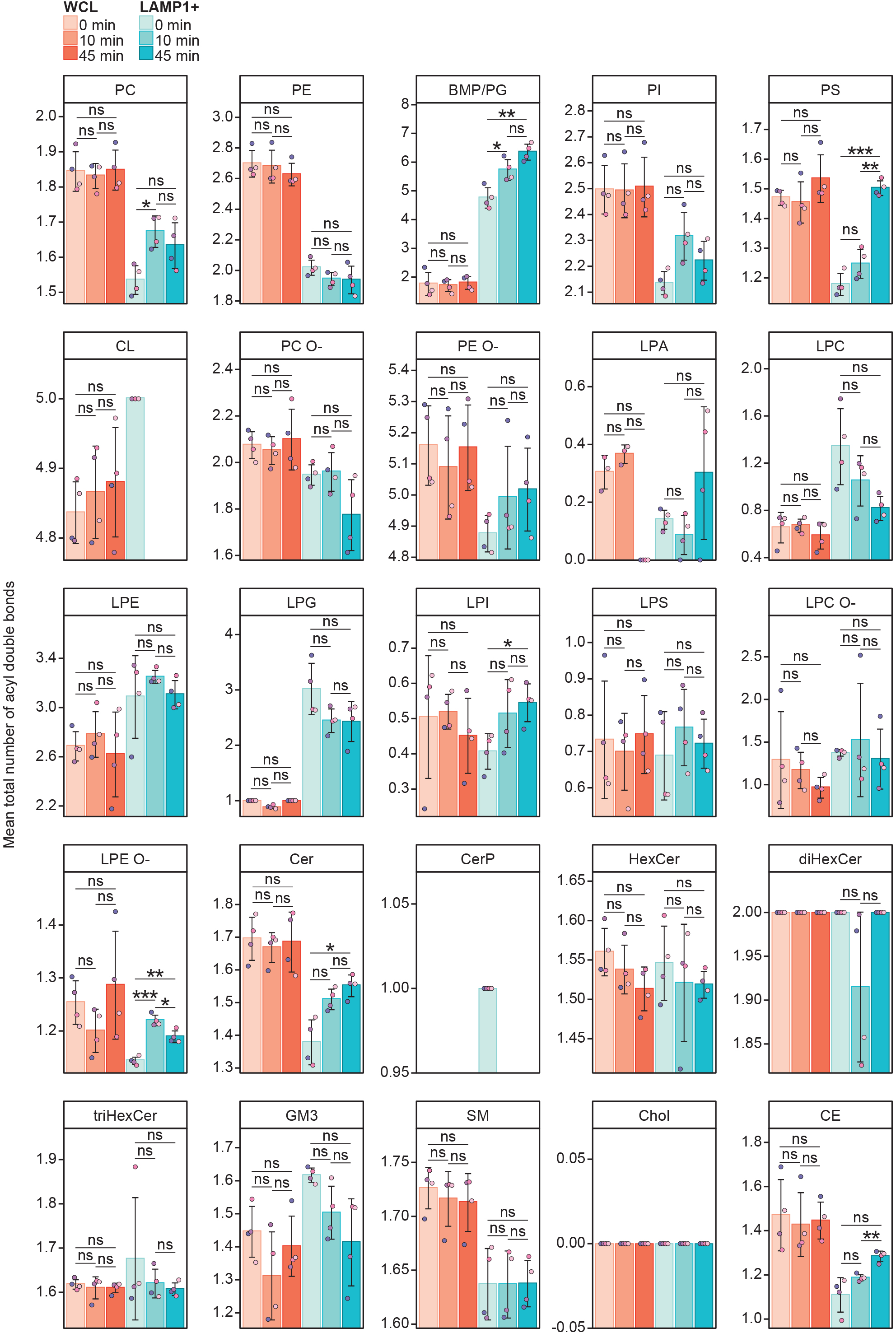
Mean total numbers of double bonds in the monitored lipid classes in the WCLs and LAMP1+ fractions from cells untreated or treated with LLOMe for 10 or 45 min. The double bonds are in the acyl (alkyl) groups of glycerophospholipids and sterol lipids, and in the acyl group and long chain base groups of sphingolipids. Multiple t-tests were performed to assess changes between means. Resultant *p*-values were corrected for multiple testing using Benjamini-Hochberg correction and differences were considered statistically significantly changed with adjusted *p*-values (q-values) of 0.05 or less. * q < 0.05; ** q < 0.01; *** q < 0.001. N=3-4. Abbreviations are as in Fig. 1.

**Figure EV4.**
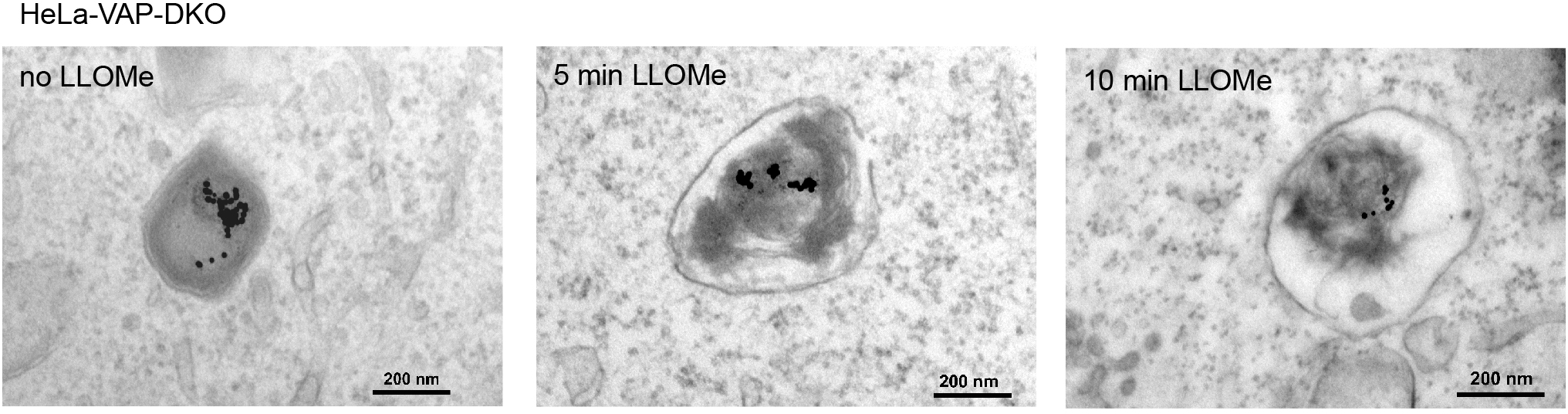
Representative electron micrographs of untreated or 250 µM LLOMe treated VAP-DKO cells. Cells were treated for 5 min or 10 min before chemical fixation. Cells were incubated with 15 nm gold nanoparticles conjugated to bovine serum albumin for 4 h to visualize lysosomes. After 4 h, gold nanoparticles were washed off and cells were incubated overnight before treatment with LLOMe and processing samples for electron microscopy. Cells treated with LLOMe show no increased membrane contacts with the endoplasmic reticulum compared to untreated cells.

**Figure EV5.**
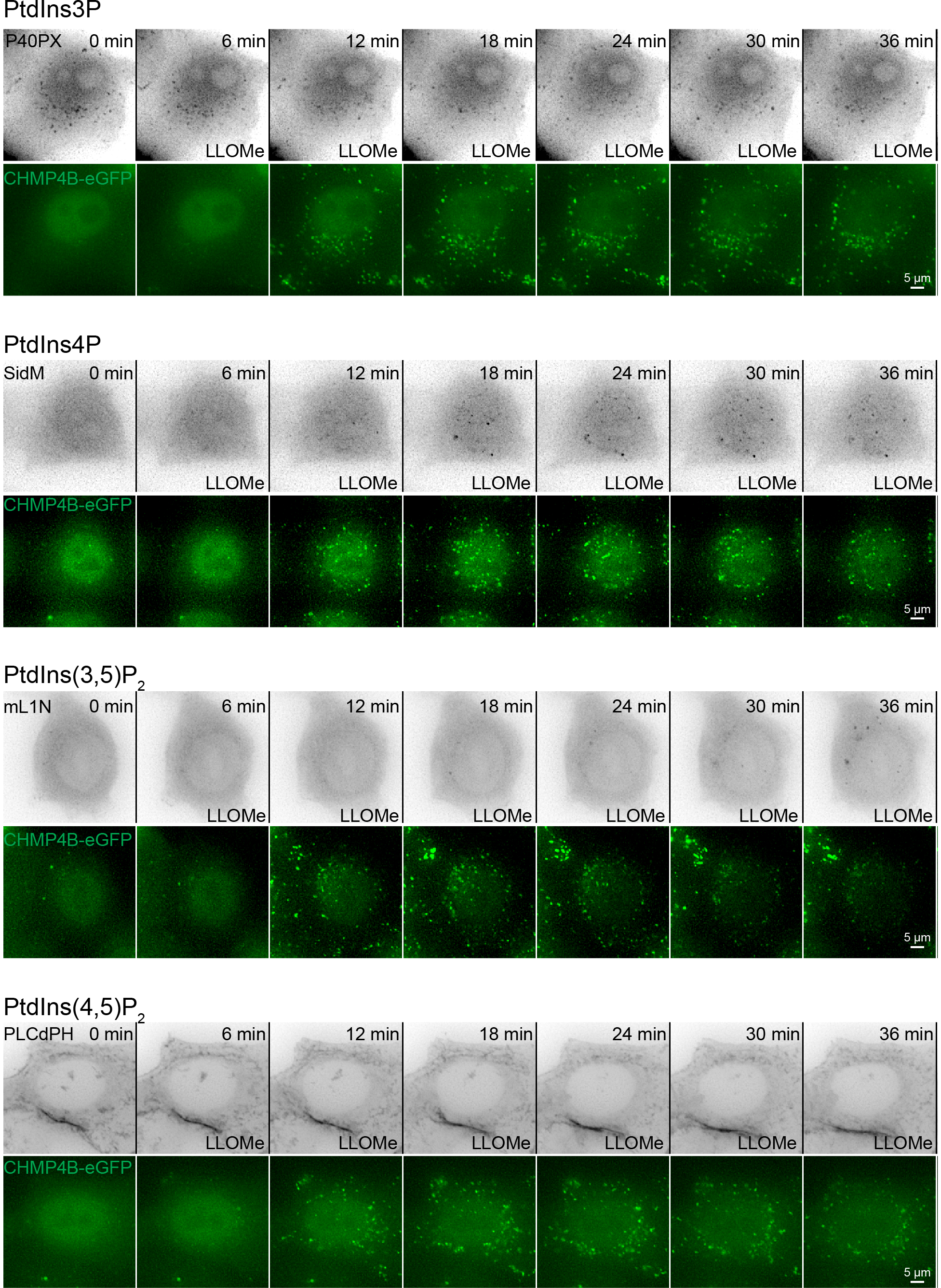
Representative movie montages of live-cell imaging experiment using mCherry tagged lipid probes in HeLa cells stably expressing CHMP4B-eGFP. P40PX was used for detection of PtdIns3P, SidM for detection of PtdIns4P, mL1N for detection of PtdIns(3,5)P_2_, and PLCdPH for detection of PtdIns(4,5)P_2_. CHMP4B-eGFP was used as a positive control of lysosomal damage. Scale bar: 5 µm.

**Figure EV6.**
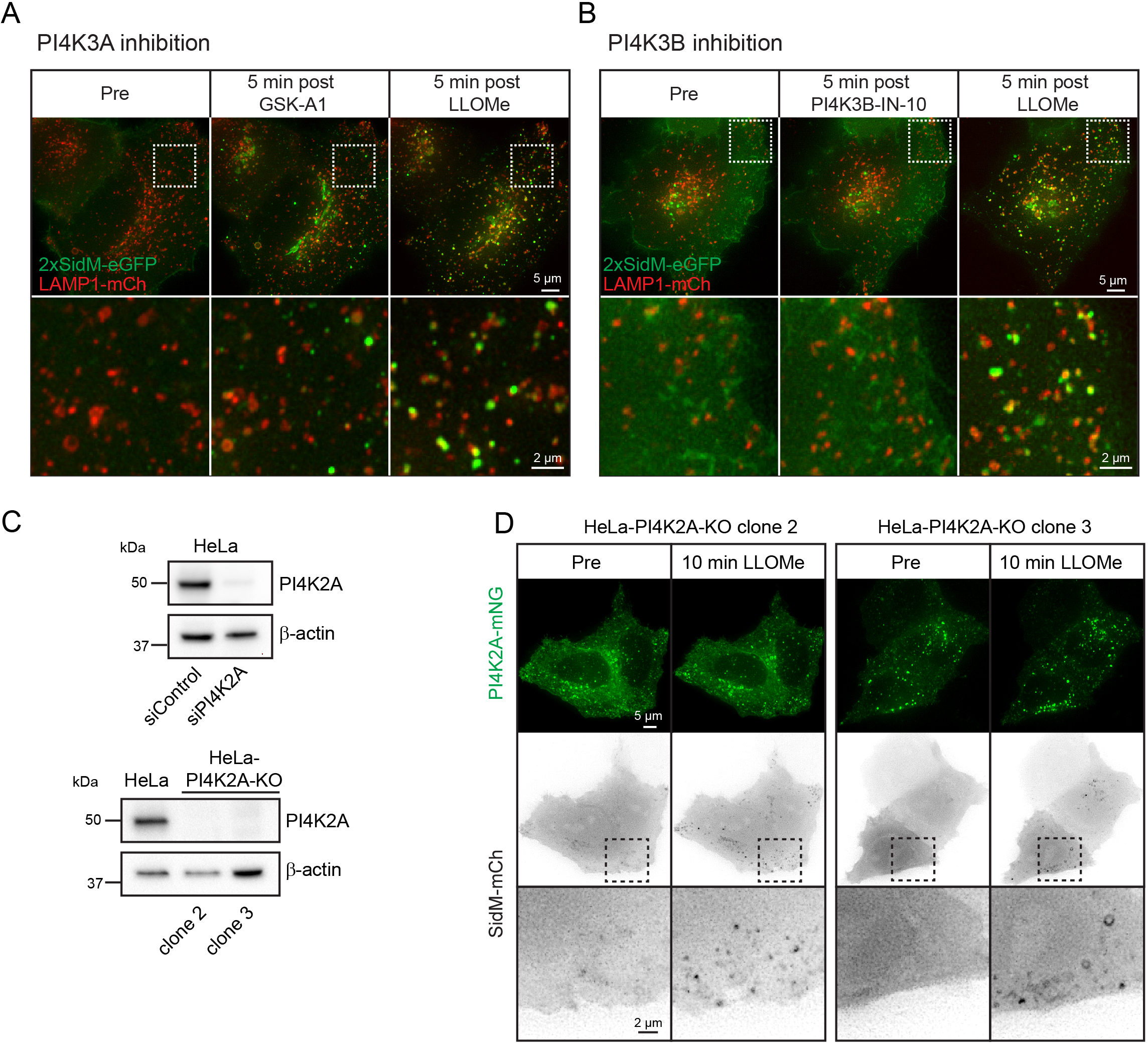
**A** Movie stills from live-cell imaging experiments of HeLa cells transiently expressing the PtdIns4P probe 2xSidM-eGFP and the lysosomal marker LAMP1-mCherry. Cells were incubated with 10 nM PI4K3A inhibitor GSK-A1 for approximately 16 min before 250 µM LLOMe was added. **B** Movie stills from live-cell imaging experiments of HeLa cells transiently expressing the PtdIns4P probe 2xSidM-eGFP and the lysosomal marker LAMP1-mCherry. Cells were incubated with 25 nM PI4K3B inhibitor PI4K3B-IN-10 for approximately 30 min before 250 µM LLOMe was added. **C** Knock down efficiency of siRNA against PI4K2A or CRISPR-Cas9 mediated knockout of PI4K2A (clone 2 and clone 3) as detected by Western blot using an anti-PI4K2A antibody. β-actin used as a loading control. **D** Exogenous expression of PI4K2A-mNG and the PtdIns4P probe SidM-mCherry in clone 2 and clone 3 of PI4K2A-KO cells shows recruitment of SidM-mCherry after incubation with 250 µM LLOMe.

**Figure EV7.**
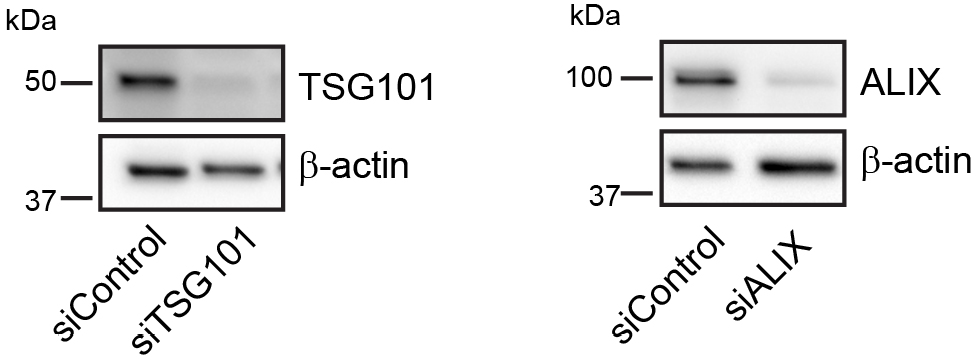
Knockdown efficiency of siRNA oligonucleotides for TSG101 or ALIX as detected by Western blot using anti-TSG101 or anti-ALIX antibodies. β-actin used as a loading control.

